# An Optimized RNF126-Targeting Covalent Handle for Molecular Glue Degraders

**DOI:** 10.64898/2026.03.06.709959

**Authors:** Aman Modi, Ethan S. Toriki, Christian E. Stieger, Emily A. Lau, Claire Song, Alyssa Chew, Amy Tsao, Kaila Nishikawa, Jeffrey McKenna, Daniel K. Nomura

## Abstract

Molecular glue degraders represent a powerful modality for targeting proteins that are refractory to traditional inhibition. However, rational design principles for molecular glue degraders remain poorly defined. Previously, we reported a chemistry-centric strategy to identify covalent degradative handles that, when appended to established ligands, convert non-degradative inhibitors into molecular glue degraders by engaging permissive E3 ligases. This effort identified a fumarate-based electrophilic handle that covalently modified the E3 ligase RNF126, enabling degradation of multiple protein targets when transplanted across diverse ligands. Despite its conceptual impact, the high intrinsic reactivity and cytotoxicity of the fumarate handle limited its translational utility. Here, we report the development of an optimized and metabolically stabilized RNF126-targeting covalent handle incorporating a *trans*-cyclobutane linker that exhibits reduced glutathione reactivity and diminished cytotoxicity while retaining robust degradative activity. When appended to the BET bromodomain inhibitor JQ1, this optimized handle yielded a potent and selective BRD4 degrader whose activity was dependent on RNF126. Importantly, transplantation of this handle onto a previously non-inhibitory ligand targeting the androgen receptor (AR) and its truncation variant, AR-V7, enabled selective degradation of both AR and AR-V7 in androgen-independent prostate cancer cells, thereby robustly inhibiting AR transcriptional activity beyond the established AR antagonist enzalutamide. Collectively, these findings demonstrate an optimized RNF126-based covalent handle for the rational development of molecular glue degraders against transcriptional regulators, including undruggable variants such as AR-V7.

## Introduction

Targeted protein degradation has emerged as a transformative approach for modulating protein function beyond the limits of classical inhibition ^1–3^. While heterobifunctional degraders such as Proteolysis Targeting Chimeras (PROTACs) have demonstrated broad utility, molecular glue degraders offer distinct advantages, including reduced molecular weight, simplified pharmacology, and the potential to engage targets lacking ligandable pockets. Despite notable clinical and preclinical successes, target-centric molecular glue discovery beyond CRBN-based degraders remains largely empirical, with limited mechanistic or chemical design rules guiding its development ^1^.

We and others have recently introduced chemistry-centric strategies for molecular glue discovery in which covalent electrophilic handles were systematically appended to known ligands to identify degradative activity ^4–8^. Using chemoproteomic profiling, we previously identified permissive E3 ligase-target pairs and demonstrated that a fumarate-based electrophile could covalently engage the E3 ligase RNF126, thereby converting otherwise non-degradative inhibitors into molecular glue degraders of their respective targets ^4^. This work provided a generalizable framework for molecular glue design by decoupling target binding from E3 ligase engagement. However, the fumarate electrophile used in this initial study exhibited high intrinsic reactivity, rapid glutathione consumption, and dose-limiting cytotoxicity, collectively constraining its translational potential. We hypothesized that refining the electrophilic geometry and linker topology of the RNF126-targeting handle would preserve productive engagement of the E3 ligase while improving metabolic stability and cellular tolerability.

Here, we report the development of a next-generation RNF126-targeting covalent handle incorporating a *trans*-cyclobutane linker that reduces electrophile reactivity while maintaining RNF126 engagement. Using BRD4 as a benchmark target, we demonstrated that this optimized handle enabled potent, selective, and RNF126-dependent degradation with minimal cytotoxicity. We further extended this strategy to androgen receptor (AR) biology, generating a molecular glue degrader capable of eliminating both full-length AR and the clinically intractable splice variant AR-V7, which lacks the ligand-binding domain and drives resistance to current therapies in prostate cancer. Together, these studies established an optimized chemical blueprint for RNF126-based molecular glue degraders and demonstrated their utility against a transcription factor previously considered undruggable.

## Results

### Optimization of an RNF126-Targeting Covalent Handle for BRD4 Degradation

We first evaluated the degradative activity and liabilities of the original fumarate-based BRD4 degrader JP-2-197 **(Figure 1a)**. Treatment of HEK293T cells with JP-2-197 led to robust degradation of both long and short BRD4 isoforms in RNF126 wild-type cells but not in RNF126 knockout cells, confirming strict RNF126 dependence **(Figure 1b-1c)**. However, JP-2-197 exhibited rapid glutathione reactivity (GSH t_1/2_ = 29 min) and pronounced cytotoxicity at higher concentrations **(Figure 1a, 1d)**, underscoring the need for a less reactive and less cytotoxic alternative.

**Figure 1.**
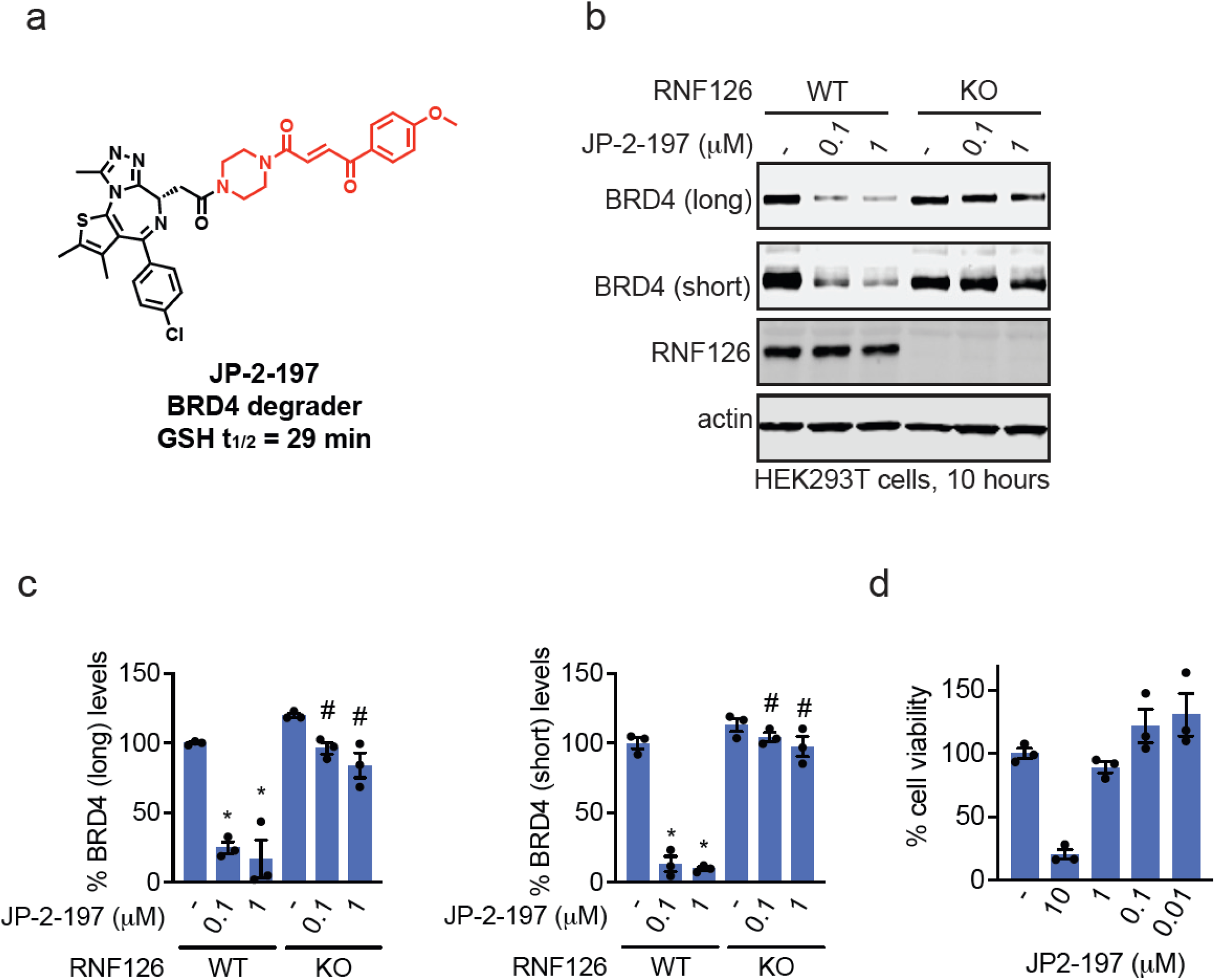
Characterization of the original fumarate-based RNF126-dependent BRD4 degrader JP-2-197. **(a)**Chemical structure of JP-2-197 and glutathione half-life. **(b)** BRD4 degradation in HEK293T cells. RNF126 wild-type (WT) and knockout (KO) cells were treated with DMSO vehicle or JP-2-197 for 10 h, after which BRD4, RNF126, and loading control actin levels were assessed by SDS/PAGE and Western blotting. **(c)** Quantification of BRD4 and RNF126 protein levels from **(c). (d)** HEK293T cell viability. HEK293T cells were treated with DMSO vehicle or JP-2-197 for 24 h, after which cell viability was assessed by CellTiter-Glo. Blot in is representative of n=3 biologically independent replicates per group. Bar graphs in **(c,d)** show individual replicates and average ± sem. Significance in **(c)** expressed as *p<0.05 compared to vehicle-treated WT control and #p<0.05 compared to respective concentrations of WT conditions.

To address these limitations, we designed a second-generation RNF126-targeting handle incorporating a *trans*-cyclobutane linker, yielding the BRD4 degrader J594 **(Figure 2a)**. This modification substantially improved glutathione stability (GSH t_1/2_ = 68 min) while preserving the covalent engagement of the pure RNF126 protein, as demonstrated by gel-based activity-based protein profiling ^9^, which showed competition for rhodamine-functionalized iodoacetamide (IA-rhodamine) labeling **(Figure 2b)**. We further demonstrated cellular RNF126 engagement and enrichment with an alkyne-functionalized version of our optimized RNF126 handle, EST2002, without enrichment of an unrelated protein, vinculin **(Figure 2c-2d)**. Importantly, J594 induced dose-dependent degradation of BRD4 with minimal effects on cell viability across the tested concentration range **(Figure 2e-2f)**.

**Figure 2.**
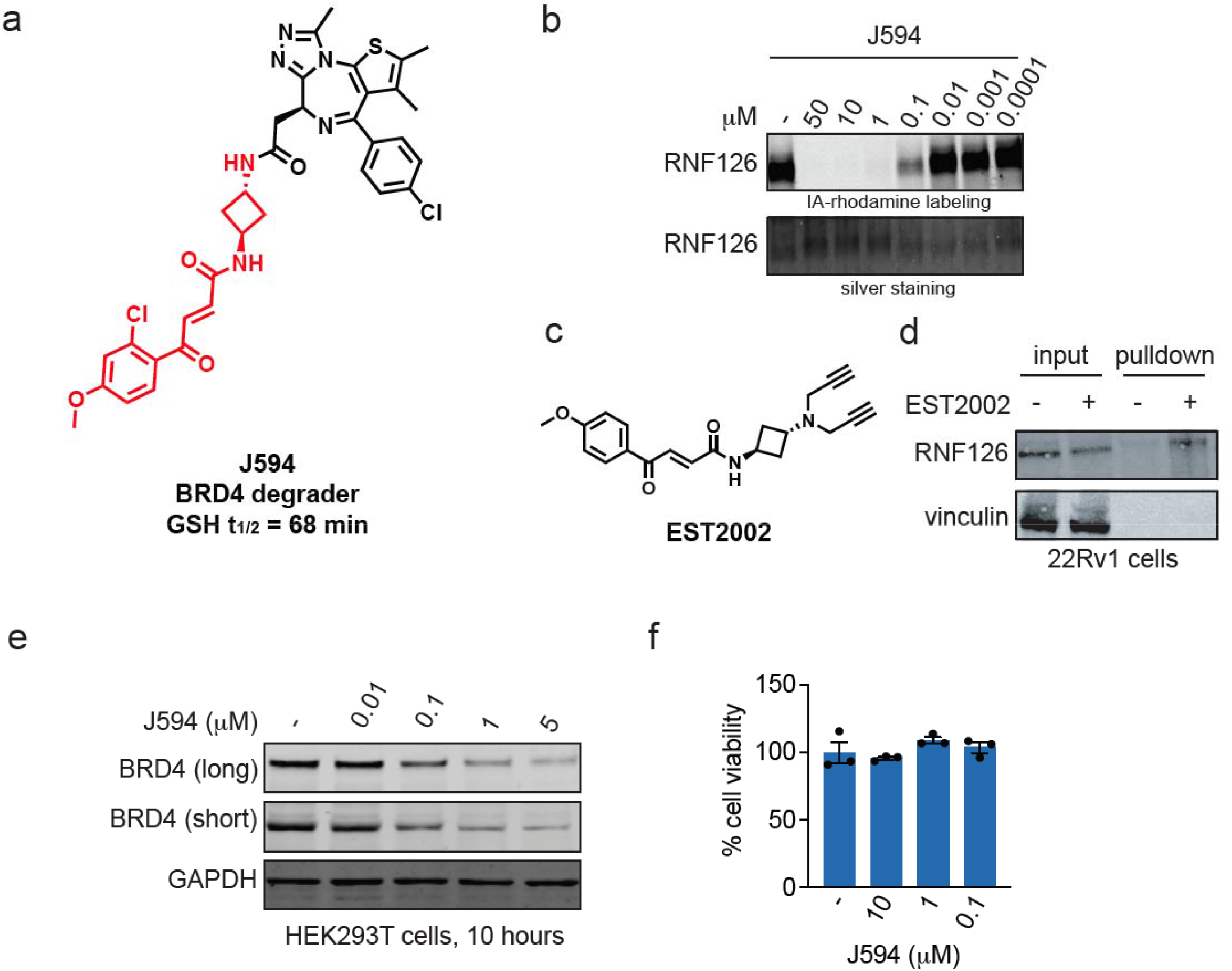
Development of an optimized RNF126-targeting BRD4 degrader. **(a)** Chemical structure of J594 with improved glutathione stability. **(b)** Gel-based ABPP of J594. RNF126 pure protein was pre-incubated with DMSO vehicle or J594 for 30 min prior to incubation with IA-rhodamine (100 nM) for 1 h, after which proteins were separated by SDS/PAGE and visualized by in-gel fluorescence. Loading was assessed by silver staining. Structure of alkyne-functionalized RNF126 handle, EST2002. **(d)** RNF126 target engagement in cells. 22Rv1 cells were treated with DMSO vehicle or EST2002 (10 µM, 2 h), after which proteomes were subjected to copper-catalyzed alkyne-azide cycloaddition (CuAAC) with a biotin-functionalized azide enrichment handle, probe-labeled proteins were avidin-enriched, and eluted. Input proteome and pulldown proteins were separated by SDS/PAGE, and RNF126 and unrelated negative control protein vinculin were visualized by Western blotting. **(e)** Dose-response of BRD4 degradation. HEK293T cells were treated with DMSO vehicle or J594 for 10 h, after which BRD4 and loading control GAPDH levels were assessed by SDS/PAGE and Western blotting. **(f)** Cell viability following J594 treatment. HEK293T cells were treated with DMSO vehicle or J594 for 24 h, after which cell viability was assessed by CellTiter-Glo. Blots in **(b,d,e)** are representative of n=3 biologically independent replicates per group. Data in **(f)** shows individual replicates and average ± sem.

We next confirmed the RNF126 dependence and BRD4 degradation selectivity mediated by J594 **(Figure 3a-3b)**. We showed that the BRD4 degradation observed upon J594 treatment in RNF126 wild-type HEK293T cells was significantly attenuated in RNF126 knockout cells **(Figure 3a-3b)**. We observed near-complete, but not complete, rescue of BRD4 degradation in knockout cells, potentially indicating that additional E3 ligases may be involved **(Figure 3a-3b)**. Quantitative proteomic analysis further demonstrated that BRD4 was the dominant protein significantly downregulated following J594 treatment, indicating high target selectivity and minimal off-target effects **(Figure 3c; Table S1)**. These results established that the optimized RNF126-targeting handle maintains the defining features of the original fumarate scaffold—potent and selective molecular glue-mediated degradation—while substantially improving metabolic and cellular tolerability.

**Figure 3.**
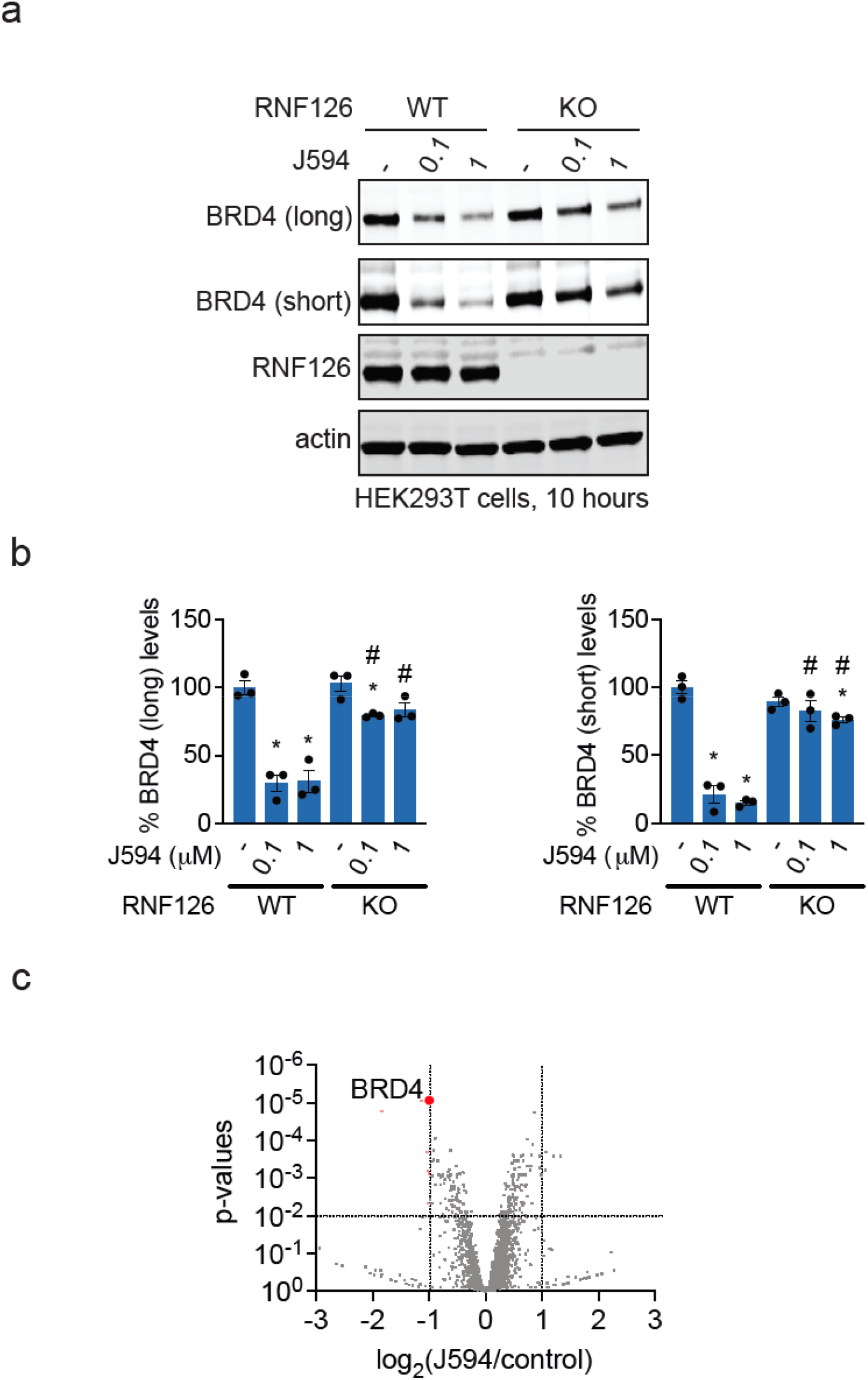
RNF126-dependent and selective BRD4 degradation by J594. **(a)** BRD4 degradation in RNF126 WT versus KO cells. RNF126 WT or KO HEK293T cells were treated with DMSO vehicle or J594 for 10 h, after which BRD4, RNF126, and loading control actin levels were assessed by SDS/PAGE and Western blotting. **(b)** Quantification of BRD4 degradation in **(a). (c)** Quantitative proteomic profiling of J594 in MBA-MB-231 cells. MDA-MB-231 cells were treated with DMSO vehicle or J594 (1 µM) for 24 h, after which proteomes were analyzed by tandem mass tagging (TMT)-based quantitative proteomics by LC-MS/MS. Blot in **(a)** is representative of n=3 biologically independent replicates per group. Bar graphs in **(b)** show individual replicates and average ± sem. Proteomic data is from n=3 biologically independent replicates per group, and the full dataset can be found in **Table S1**. Significance in **(b)** is expressed as *p<0.05 compared to vehicle-treated WT control and #p<0.05 compared to respective concentrations of WT conditions.

### Transplantation of the Optimized Handle Enables Degradation of AR and AR-V7

Having validated the optimized handle through observing BRD4 degradation, we next sought to test whether this handle could be transplanted onto a ligand targeting the AR and its splice variant AR-V7. AR-V7 lacks the ligand-binding domain targeted by approved AR antagonists and is a key driver of resistance in androgen-independent prostate cancer ^10,11^. Although AR antagonists and AR PROTACs are in clinical use or in clinical development ^12,13^, these molecules all target the ligand-binding domain and are ineffective against AR-V7. AR-V7 is considered undruggable since the remaining N-terminal and DNA-binding domains are largely unstructured and intrinsically disordered ^10,11^. While our lab has previously identified a direct-acting, selective, covalent destabilizing degrader that targets the intrinsically disordered C125 on AR and AR-V7, this degrader is not potent and is stoichiometric due to its covalent engagement with the target protein^14^. A molecular glue degrader with reversible AR-V7 targeting would be more desirable for enabling sub-stoichiometric and catalytic degradation of the target ^2^.

We synthesized the AR/AR-V7 degrader EST1140 by appending the optimized RNF126-targeting handle to VPC-14228 **(Figure 4a)** ^15,16^. While this ligand has been previously shown to bind to the DNA-binding domain of AR and act as a functional antagonist of AR transcriptional activity, promoting its usein the development of PROTACs, in our hands, VPC-14228 does not inhibit, but instead stimulates AR transcriptional activity in 22Rv1 prostate cancer cells expressing AR and AR-V7 **(Figure S1a)**. We have also shown that previously disclosed PROTACs using this ligand do not degrade AR or AR-V7 ^4^. In contrast, we demonstrated that our original RNF126-based fumarate handle was capable of degrading AR and AR-V7, although it was also cytotoxic and did not exhibit the desired selectivity ^4^. Nonetheless, to further confirm that VPC-14228 engages AR and AR-V7 in cells, we synthesized an alkyne-functionalized photoaffinity probe based on VPC-14228 and demonstrated that this probe could be used to engage and enrich AR and AR-V7 in 22Rv1 cells **(Figure S1c-S1d)**. We appended our optimized fumarate handle onto VPC-14228 to generate EST1140. We demonstrated that EST1140 still covalently bound to pure RNF126 protein, as demonstrated by competition with IA-rhodamine labeling **(Figure 4b)**. EST1140 treatment in 22Rv1 prostate cancer cells, which express both AR and AR-V7, led to a dose- and proteasome-dependent degradation of both proteins **(Figure 4c-4d)**. We further demonstrate that the covalent handle was necessary for this degradation as a non-reactive derivative AM-2-040 was incapable of degrading AR and AR-V7 **(Figure S1e-S1f)**. These data contrasted with the clinically approved prostate cancer AR antagonist enzalutamide, which does not alter AR or AR-V7 levels, and with the ARV110 PROTAC under clinical development, which eliminates only full-length AR, not AR-V7 **(Figure 4e)**. Quantitative proteomics revealed highly selective degradation of AR/AR-V7 with minimal off-target effects **(Figure 4f; Table S2)**. Importantly, EST1140-mediated degradation of FLAG-tagged AR was abrogated in RNF126 knockout cells, establishing RNF126 dependence **(Figure 4g-4h)**.

**Figure 4.**
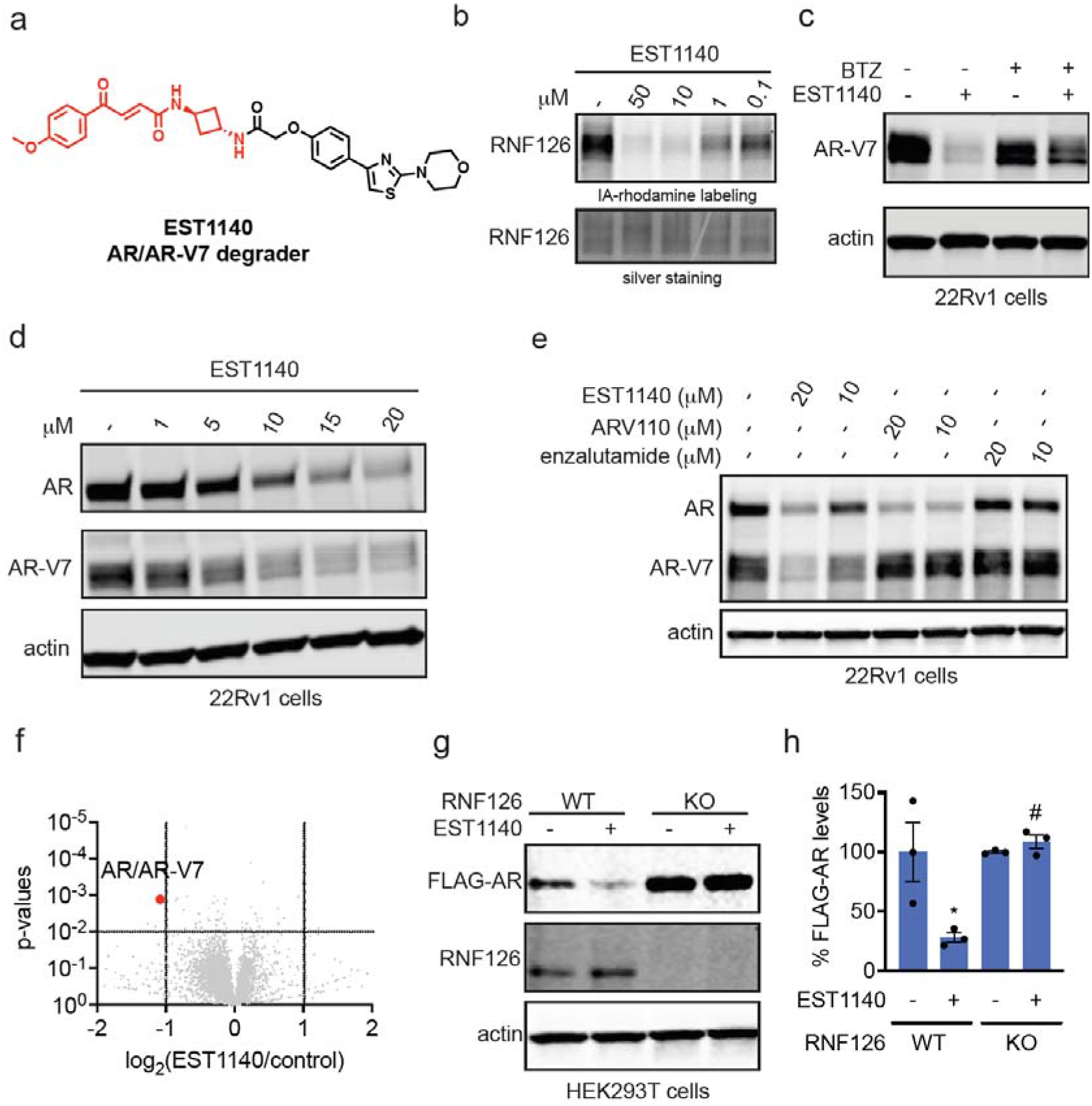
Transplantation of the optimized RNF126 handle enables AR and AR-V7 degradation. **(a)** Chemical structure of EST1140. **(b)** Gel-based ABPP of EST1140. RNF126 pure protein was pre-incubated with DMSO vehicle or EST1140 for 30 min prior to incubation with IA-rhodamine (100 nM) for 1 h, after which proteins were separated by SDS/PAGE and visualized by in-gel fluorescence. Loading was assessed by silver staining. **(c)** Proteasome-dependence of AR-V7 degradation. 22Rv1 cells were pre-treated with DMSO vehicle or proteasome inhibitor bortezomib for 1 h prior to treatment of cells with DMSO vehicle or EST1140 (20 µM) for 24 h, after which AR-V7 and loading control actin levels were assessed by SDS/PAGE and Western blotting. **(d**,**e)** Dose-response of EST1140, ARV110, and enzalutamide. 22Rv1 cells were treated with DMSO vehicle, EST1140, ARV110, or enzalutamide for 24 h, after which AR, AR-V7, and loading control actin levels were assessed by SDS/PAGE and Western blotting. **(f)** Quantitative proteomic profiling of EST1140 in 22Rv1 cells. 22Rv1 cells were treated with DMSO vehicle or EST1140 (20 µM) for 24 h, after which proteomes were analyzed using TMT-based quantitative proteomics by LC-MS/MS. **(g)** AR degradation. FLAG-AR-expressing RNF126 WT versus KO HEK293T cells were treated with DMSO vehicle or EST1140 (20 µM) for 24 h, after which FLAG-AR, RNF126, and loading control actin levels were assessed by SDS/PAGE and Western blotting. **(h)** Quantification of the experiment described in **(h)**. Blots in **(b-e, g)** are representative of n=3 biologically independent replicates per group. Proteomic data in **(f)** is from n=3 biologically independent replicates per group, and the full dataset can be found in **Table S2**. Bar graph in **(h)** shows individual replicates and average ± sem. Significance in **(h)** is expressed as *p<0.05 compared to vehicle-treated WT control and #p<0.05 compared to EST1140-treated WT groups.

### EST1140 Disrupts AR and AR-V7 Stability and Suppresses AR Transcriptional Activity

To further assess AR and AR-V7 target engagement by EST1140 and the functional consequences of AR/AR-V7 degradation, we performed cellular thermal shift assays (CETSA) in 22Rv1 cells ^17^. Interestingly, while EST1140 does not covalently engage AR or AR-V7, EST1140 treatment significantly destabilized both AR and AR-V7, resulting in marked reductions in thermal stability relative to control conditions **(Figure 5a-5b)**. First, this CETSA-based destabilization confirmed that EST1140 engaged AR and AR-V7 in 22Rv1 cells. Second, these data suggest that EST1140 degrades these proteins not only via RNF126-dependent molecular glue degrader mechanisms but also through the destabilization of AR and AR-V7. These data were reminiscent of our recently disclosed covalent destabilizing degrader of AR and AR-V7, EN1441, which acts by covalently modifying C125 on AR/AR-V7 ^14^. Functionally, EST1140 robustly inhibited AR-dependent transcriptional activity with an EC_50_ of approximately 8 µM, compared to enzalutamide, which only showed a partial inhibition of total AR transcriptional activity in 22Rv1 cells, with less than 50 % inhibition at even the highest concentrations, since enzalutamide does not bind to AR-V7 **(Figure 5c-5d)**. These findings demonstrated that RNF126-based molecular glue degradation provides a distinct and more effective mechanism for suppressing AR transcriptional activity in androgen-independent prostate cancer cells.

**Figure 5.**
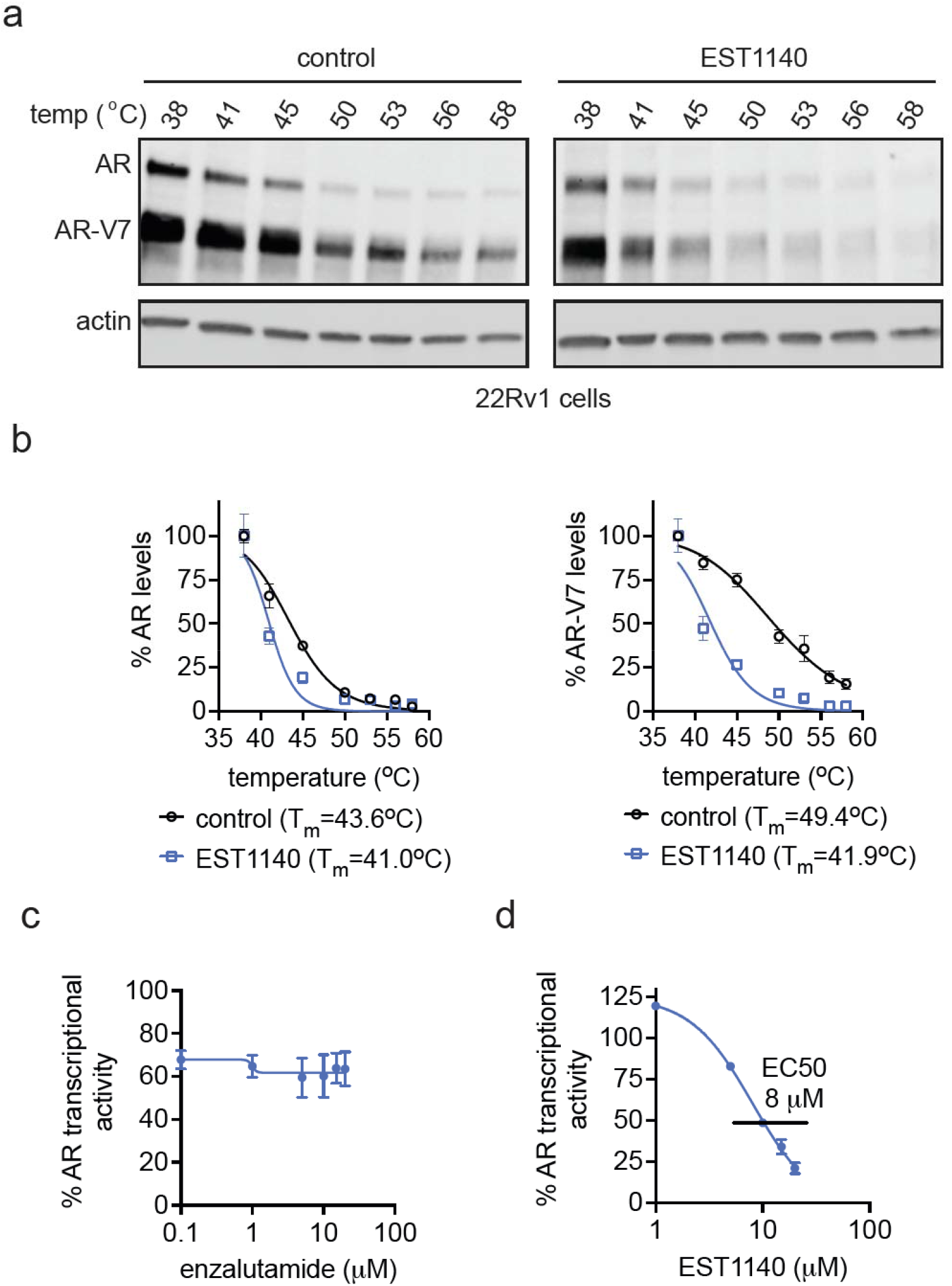
Functional consequences of AR/AR-V7 degradation by EST1140. **(a)** CETSA analysis demonstrating destabilization of AR and AR-V7. 22Rv1 cells were treated with DMSO vehicle or EST1140 (10 µM) for 1 h, after which cells were heated to designated temperatures, precipitated proteins were cleared, and AR, AR-V7, and actin control levels were assessed by SDS/PAGE and Western blotting. **(b)** Quantification of CETSA data in **(a). (c**,**d)** Inhibition of AR transcriptional activity by EST1140. 22Rv1 cells expressing an AR luciferase reporter were treated with DMSO vehicle, enzalutamide **(c)**, or EST1140 **(d)** for 12 h, and total AR/AR-V7 transcriptional activity was assessed by luminescence. Blot in **(a)** is representative of n=3 biologically independent replicates per group. Graphs in **(b-d)** show average ± sem values of n=3 biologically independent replicates per group.

## Discussion

This study advances molecular glue degrader design by demonstrating that the liabilities of highly reactive covalent handles can be mitigated through chemical optimization without sacrificing degradative potency or selectivity. By refining the geometry and linker composition of a fumarate-based RNF126-targeting electrophile, we developed a second-generation covalent handle with improved glutathione stability, reduced cytotoxicity, and preserved E3 ligase engagement.

Using BRD4 as a benchmark, we showed that this optimized handle supports potent, selective, and RNF126-dependent degradation with minimal off-target effects. Importantly, the modularity of this handle enabled its transplantation onto a previously non-functional AR-V7 ligand, yielding a molecular glue degrader capable of eliminating both full-length AR and the classically undruggable AR-V7 variant. This capability addresses a major unmet need in prostate cancer therapy, where resistance driven by AR-V7 remains a significant challenge.

The work described here builds on our and others’ prior efforts to establish a chemistry-centric framework for the rational design of molecular glue degraders via covalent engagement of E3 ligases ^4–8^. In addition to RNF126, we have previously identified multiple covalent degradative handles that act through DCAF16 by selectively targeting C119 or a second cysteine cluster encompassing C173/C178 ^5,6^. These studies demonstrated that molecular glue degradation can be achieved by engaging distinct cysteine sites within the same E3 ligase and that productive degradation is governed not simply by covalent binding per se, but by the precise spatial and chemical compatibility between the electrophilic handle, the E3 ligase surface, and the recruited target protein. Work from the Gray, Fischer, and Winter laboratories further established the importance of DCAF16 as a permissive E3 ligase for covalent molecular glue degraders, particularly through engagement of C58, in the context of BRD4 degradation, in addition to highlighting the significance of additional E3 ligases, including DCAF11 and FBXO22 ^7,8,18,19^. Collectively, these studies also extend the original work by Cravatt and Zhang on covalent targeting of DCAF16 and work by our lab in targeting RNF114 and RNF4 using electrophilic handles to enable targeted protein degradation ^9,20,21^.

Importantly, the convergence of these independent efforts highlights a unifying principle for molecular glue degrader discovery: covalent engagement of E3 ligase cysteines represents a powerful and generalizable strategy for enabling degradation, but success hinges on treating the electrophilic handle as an optimizable chemical module rather than a fixed warhead. By demonstrating that an optimized RNF126-targeting handle can be transplanted across structurally and biologically distinct targets—including the transcription factor variant AR-V7—this work extends the scope of covalent molecular glue degraders beyond classically druggable targets such as BET proteins and kinases. It reinforces the value of a modular, handle-centric design paradigm.

Future efforts will focus on expanding the scope of RNF126-based molecular glues, further tuning electrophile reactivity for in vivo applications, and exploring the generalizability of this approach across additional E3 ligases and disease-relevant targets. Beyond RNF126, RNF4, DCAF16, FBXO22, DCAF11, and FEM1B ^5,6,8,9,18,20–26^, future efforts will focus on identifying new covalent degradative handles and permissive E3 ligase pairs that can be harnessed for the rational design of molecular glue degraders. Taken together, this work provides both a practical molecular glue scaffold and a conceptual framework for advancing covalent molecular glue degraders toward therapeutic development.

## Acknowledgment

We thank the members of the Nomura Research Group and Novartis BioMedical Research for critically reading the manuscript. This work was also supported by Novartis Biomedical Research, the National Science Foundation Molecular Foundations for Biotechnology (MFB) grant (2127788), the UC Berkeley Molecular Therapeutics Initiative (MTI), Bakar Fellows Award, the Mark Foundation for Cancer Research ASPIRE Award, and the National Institutes of Health (R35CA263814, R01CA240981, UM1CA29410). We also thank Hasan, Lund, and the UC Berkeley NMR facility in the College of Chemistry (CoC-NMR) and Zhongrui at the QB3/chemistry mass spectrometry facility for spectroscopic assistance. Instruments in the College of Chemistry NMR facility are partly supported by NIH S10OD024998. K. L. acknowledges the National Science Foundation for a pre-doctoral graduate fellowship. RNA Sequencing was performed at the QB3 Genomics Facility at UC Berkeley, Berkeley, CA RRID:SCR_022170 and was supported by NIH S10 OD018174 Instrumentation Grant.

## Author Contributions

AM, EST, DKN conceived of the project. AM, EST, DKN, JM performed experiments, analyzed data, and interpreted results. AM, JM, MS, and DKN wrote the paper.

