## Supporting Information for "An Optimized RNF126-Targeting Covalent Handle for Molecular Glue Degraders"

### Table Legends

**Table S1. Quantitative proteomic profiling of J594 in MBA-MB-231 cells.** MDA-MB-231 cells were treated with DMSO vehicle or J594 (1 μM) for 24 h, after which proteomes were analyzed by tandem mass tagging (TMT)-based quantitative proteomics by LC-MS/MS. Data are from n=3 biologically independent replicates per group.

**Table S2. Quantitative proteomic profiling of EST1140 in 22Rv1 cells.** 22Rv1 cells were treated with DMSO vehicle or EST1140 (20 μM) for 24 h, after which proteomes were analyzed using TMT-based quantitative proteomics by LC-MS/MS. Data are from n=3 biologically independent replicates per group.

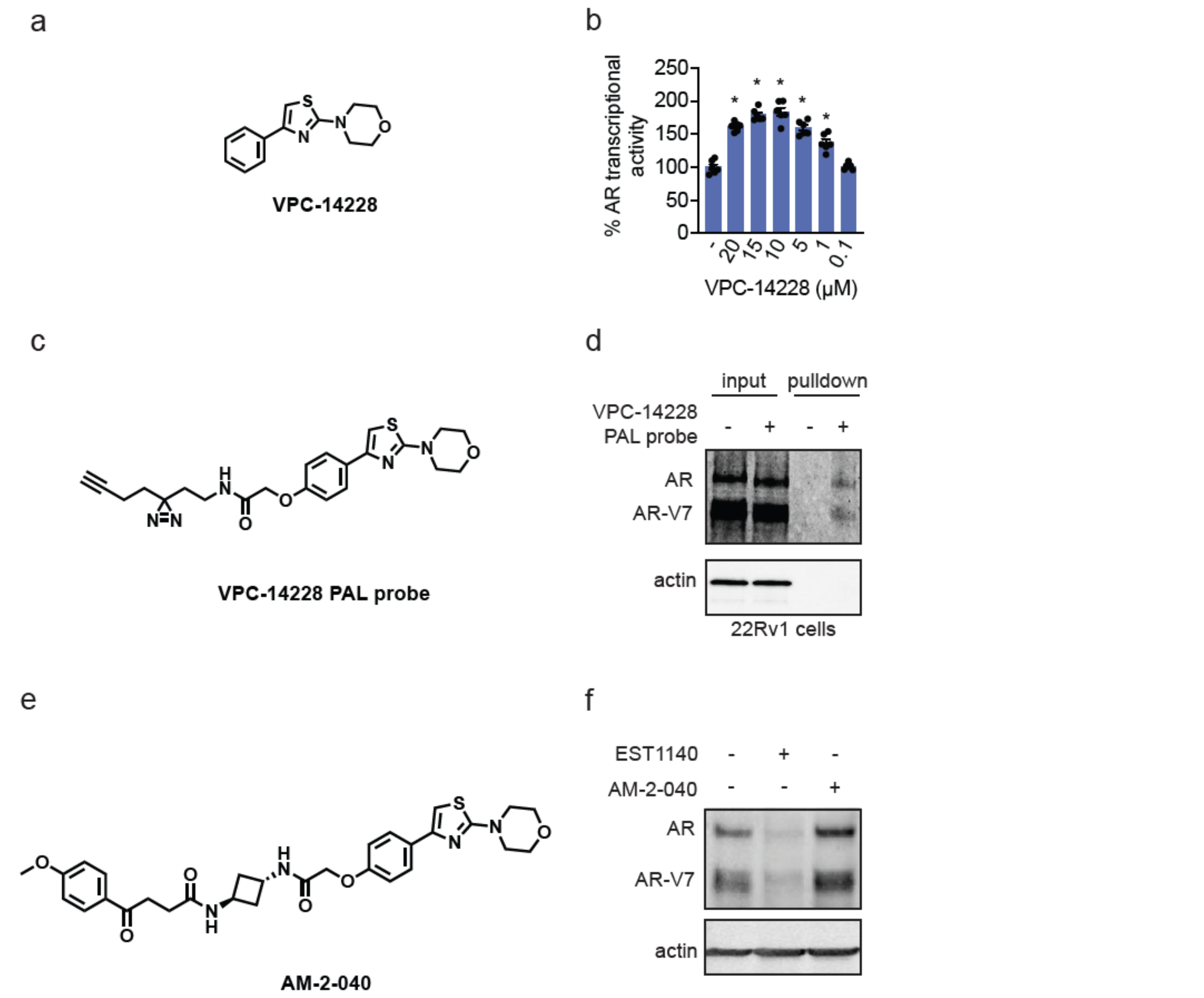


**Figure S1. Characterization of VPC-14228 and non-reactive analogs of EST1140. (a)** Structure of VPC-14228. **(b)** AR transcriptional activity in 22Rv1 cells. 22Rv1 cells expressing an AR luciferase reporter were treated with DMSO vehicle or VPC-14228 for 12 h, and total AR/AR-V7 transcriptional activity was assessed by luminescence. **(c)** Structure of alkyne-functionalized photoaffinity (PAL) probe based on VPC-14228. **(d)** Photoaffinity pulldown of AR and AR-V7. 22Rv1 cells were treated with DMSO vehicle or VPC-14228 PAL probe (20 μM) for 1h, after which cells were irradiated with hν for 30 min. Proteomes were subjected to copper-catalyzed alkyne-azide cycloaddition (CuAAC) with a biotin-functionalized azide enrichment handle, probe-labeled proteins were avidin-enriched, and eluted. Input proteome and pulldown proteins were separated by SDS/PAGE, and AR, AR-V7, and unrelated negative control protein actin were visualized by Western blotting. **(e)** Structure of unreactive analog of EST1140, AM-2-040. **(f)** Comparing AR and AR-V7 degradation with EST1140 and AM-2-040. 22Rv1 cells were treated with DMSO vehicle, EST1140 (20 mM), or AM-2-040 (20 mM) for 24 h, after which AR, AR-V7, and loading control actin levels were assessed by SDS/PAGE and Western blotting. Bar graph in **(b)** show individual replicates and average ± sem from n=5 biologically independent replicates per group. Blots in **(d,f)** are representative of n=3 biologically independent replicates per group. Significance in **(b)** shown as *p<0.05 compared to vehicle-treated controls.

### Supporting Methods

#### Cell Culture

HEK293T and MDA-MB-231 cells were purchased from the UC Berkeley Cell Culture Facility and were cultured in DMEM containing 10% (v/v) FBS. 22Rv1 cells were purchased from the UC Berkeley Cell Culture Facility and were cultured in RPMI 1640 containing 10% (v/v) FBS. AIZ-AR stable 22Rv1 luciferase reporter cells were purchased from Applied Biological Materials (T3104) and were cultured in RPMI 1640. All cell lines were maintained at 37°C with 5% $CO_2$.

#### Expression and purification of recombinant RNF126 protein

RNF126 mammalian expression plasmid with a C-terminal FLAG tag was purchased from Origene (Origene Technologies Inc., RC204986). The plasmid was transformed into NEB 5-alpha Competent E. coli (DH5$\alpha$) cells (NEB product no. C2987H). The following day, a single transformed colony was used to inoculate 50 mL of nutrient rich LB medium containing kanamycin (50 µg/mL) and was incubated at 37 °C overnight, with agitation (250 rpm). A Miniprep (Qiagen) kit was used to isolate the plasmid before sequence verification with appropriate primers. HEK293T cells were grown to 30-50% confluency in DMEM supplemented with 10% FBS (Corning) and maintained at 37°C with 5% CO2. Immediately before transfection, media was replaced with DMEM containing 10% FBS. Each plate was transfected with 24 µg of overexpression plasmid with 24 µL Lipofectamine 2000 (Invitrogen) in Opti-MEM. After 48 h cells were collected in PBS, lysed by sonication, and batch bound with anti-DYKDDDDK resin (GenScript, L00432) for 2 hours. Lysate and resin were washed with PBS and eluted with 133.33 µg/mL 3X FLAG peptide (ApexBio, A6001) in PBS. Five elutions were performed for 15 minutes each. Elutions were concentrated and the protein was stored in PBS. Concentration and purity were determined using the BCA assay and Western blotting.

#### Cell Lysate Preparation

Cells were washed with PBS, harvested by scraping, and pelleted by centrifugation (1400*g*, 4 min, 4 °C). Pellets were resuspended in PBS, and lysed by sonication or RIPA lysis buffer (supplemented with Pierce protease inhibitor mini tablets, EDTA-free, Thermo Fisher Scientific, A32955). Unless otherwise stated, total lysate was utilized for Western blotting and chemoproteomics experiments. Protein concentrations were determined using the BCA assay (Pierce, 23225). Lysates were stored at −80°C, or taken for Western blotting or chemoproteomics experiments.

#### Western Blotting

Anti-GAPDH (Cell Signaling Technology, D4C6R), β-actin (Cell Signaling Technology, E4D9Z), BRD4 (Abcam ab128874), AR (Cell Signaling Technology, D6F11), RNF126 (Santa Cruz Biotechnology sc-376005), were obtained, and dilutions were prepared in accordance with the manufacturer's recommended protocol. Proteins were resolved by SDS/PAGE and transferred to nitrocellulose membranes using the Trans-Blot Turbo transfer system (Bio-Rad). Membranes were blocked with 5% BSA in Tris-buffered saline containing Tween 20 (TBS-T) solution for 1 hr at RT washed in TBS-T, and probed with primary antibody diluted in recommended diluent per manufacturer overnight at 4 °C. After 3 washes with TBS-T, the membranes were incubated in the dark with IR680- or IR800-conjugated secondary antibodies at 1:10,000 dilution in 5 % BSA in TBS-T at RT for 1 h. After 3 additional washes with TBST, blots were visualized using an Odyssey Li-Cor fluorescent scanner. The membranes were stripped using ReBlot Plus Strong Antibody Stripping Solution (EMD Millipore, 2504) when additional primary antibody incubations were performed.

#### Knockdown Studies

Short-hairpin oligonucleotides were used to knock down the expression of RNF126 in HEK293T cells. For lentivirus production, lentiviral plasmids and packaging plasmids (pMD2.G, Addgene catalog no. 12259 and psPAX2, Addgene catalog no. 12260) were transfected into HEK293T cells using Lipofectamine 2000 (Invitrogen). Lentivirus was collected from filtered cultured medium and used to infect the target cell line with 1:1000 dilution of polybrene. Target cells were selected over 3 days with 1 µg/mL of puromycin for HEK293T cells. The short-hairpin sequences which were used for generation of the knockdown lines were: RNF126: TGCCATCATCACACAGCTCCT (Sigma RNF126 MISSION shRNA Bacterial Glycerol Stock, TRCN0000368954). MISSION TRC1.5 pLKO.1- or TRC2 pLKO.5-puro Non-Mammalian shRNA Control (Sigma) was used as a control shRNA.

**Gel Based ABPP**

Recombinant RNF126 (0.1µg/sample) was pre-treated with either DMSO vehicle or covalent ligand at 37 °C for 30 min in 25 µL of PBS and subsequently treated with IA-Rhodamine (concentrations designated in figure legends) (Setareh Biotech) at room temperature for 1 h in the dark. The reaction was stopped by addition of 4x reducing Laemmli SDS sample loading buffer (Alfa Aesar). After boiling at 95 °C for 5 min, the samples were separated on precast 4−20% Criterion TGX gels (Bio-Rad). Probe-labeled proteins were analyzed by in-gel fluorescence using a ChemiDoc MP (Bio-Rad).

**Quantitative TMT Proteomics Analysis**

Cells were treated with DMSO vehicle or compound for 24 h and lysates were prepared as described above. 25-100ug of lysate was reduced, alkylated and digested with sequencing grade trypsin overnight. Individual samples were then labeled with isobaric tags using commercially available TMTsixplex (Thermo Fisher Scientific, P/N 90061) kits, in accordance with the manufacturer's protocols. Tagged samples (20 µg per sample) were combined, dried using a vacuum concentrator at 30 °C, resuspended with 300 µL 0.1% TFA in $H_2O$, and fractionated using high pH reversed-phase peptide fractionation kits (Thermo Fisher Scientific, P/N 84868) according to the manufacturer's protocol. Fractions were dried using a vacuum concentrator at 30 °C, resuspended with 50 µL 0.1% FA in $H_2O$, and analyzed by LC-MS/MS as described below.

Quantitative TMT-based proteomic analysis was performed as previously described using a Thermo Eclipse with FAIMS LC-MS/MS. Acquired MS data was processed using ProLuCID (in IP2 v.3-v.5, (Integrated Proteomics Applications, Inc.)) or SagePro (in Chaparral Laboratories, Inc.). Trypsin cleavage specificity (cleavage at K, R except if followed by P) allowed for up to 2 missed cleavages. Carbamidomethylation of cysteine was set as a fixed modification, methionine oxidation and TMT-modification of N-termini and lysine residues were set as variable modifications. Reporter ion ratio calculations were performed using summed abundances with most confident centroid selected from a 20 ppm window. Only peptide-to-spectrum matches that are unique assignments to a given identified protein within the total dataset are considered for protein quantitation. High confidence protein identifications were reported with a <1% false discovery rate (FDR) cut-off. Differential abundance significance was estimated using ANOVA with Benjamini-Hochberg correction to determine p-values.

**AM-2-012 Pure Protein Competitive Click Pulldown**

0.5 µg of RNF126 recombinant protein was treated with DMSO vehicle or 50µM EST1140 in 50 µL PBS for 30 min at 37°C. Samples were subsequently treated with DMSO vehicle or AM-2-012 for 30 min at RT. CuAAC) was performed by addition of 0.25 µL Azide-Fluor 545 (5mM in DMSO, Click Chemistry Tools, Inc. product #AZ109-5), 1 µL $CuSO_4$ (50mM in $H_2O$), 3 µL TBTA (0.9 mg/mL in 4:1 tBuOH/DMSO) and 1 µL TCEP (14.4 mg/mL in $H_2O$). After one hour at RT, 30 µL 4x Laemlli's buffer was added. Probe-labeled proteins were analyzed by in-gel fluorescence using a ChemiDoc MP (Bio-Rad).

**EST2002 pulldown method?**
**Cellular Thermal Shift Assay (CETSA)**

22Rv1 cells were harvested by scraping from a 10 cm dish after 1 hour at 50 µM compound treatment. Cells were resuspended in PBS containing protease inhibitor cocktail and 50 µM compound and then aliquoted into eight 0.2 mL PCR strips with 100 µL per tube. PCR strips were designated a temperature via a gradient program on Bio-Rad's T100 Thermal cycler (Bio-Rad, 1861096). Samples were heated at their respective temperatures (37°C, 38°C, 41°C, 45°C, 50°C, 53°C, 56°C, 58°C) for 3 minutes and then at 25 °C for 3 minutes. Immediately following, cells were snap-lysed using liquid nitrogen (3 freeze-thaw cycles). Cell debris, along with any precipitated and aggregated proteins, were removed by centrifugation at 20,000 g for 20 minutes at 4

°C. 80 µL of supernatant was transferred to new PCR strips, 26.6 µL of 4x reducing Laemmli SDS sample loading buffer was added to each sample and boiled at 95°C for 10 minutes. Probe-labeled proteins were analyzed by in-gel fluorescence using an Odyssey DLx Imager (LICORbio).

### General Procedures

#### Amide Couplings

#### General Procedure A

A mixture of the carboxylic acid (1.1 equiv.) and HATU (2 equiv.) was purged with $N_2$ for 5 minutes. The mixture was dissolved in *N,N*-dimethylformamide (0.1M). *N,N*-diisopropylethylamine (DIPEA, 3 equiv.) was transferred to the reaction mixture. The reaction was stirred at ambient temperature for 15 minutes. The amine was purged with $N_2$ in a separate vial, dissolved in a minimal amount of DMF and transferred to the reaction. The reaction mixture was stirred at ambient temperature overnight. The reaction was quenched with 5X volume of 5% LiCl (aq.) and extracted thrice with ethyl acetate (EtOAc) or dichloromethane (DCM). Organic layers were dried over $Na_2SO_4$, filtered and concentrated *in vacuo*. Crude reactions were purified by silica gel flash chromatography to yield the title compound.

#### General Procedure B

The carboxylic acid (1.1 equiv.) was purged with $N_2$ for 5 minutes and dissolved in DMF (0.1M). DIPEA (3 equiv.) to which propylphosphonic anhydride solution (>50 wt.% in EtOAc, 1.1 equiv.) was transferred. The reaction mixture was stirred at ambient temperature for 30 minutes. The amine was purged with $N_2$ in a separate vial, dissolved in a minimal amount of DMF and transferred. The reaction was stirred at ambient temperature overnight. The reaction was quenched with 5X volume of 5% LiCl (aq) and extracted thrice with ethyl acetate (EtOAc) or dichloromethane (DCM). Organic layers were dried over $Na_2SO_4$, filtered and concentrated *in vacuo*. Crude reactions were purified by silica gel flash chromatography to yield the title compound.

#### General Procedure C

The corresponding tert-butyloxycarbonyl protected amine was dissolved in a 1:1 mixture of DCM:Trifluoroacetic acid (30 equiv.). The reaction mixture was stirred for 30 min – 1 hour. The volatiles were removed *in vacuo* and crude residue was used without further purification.

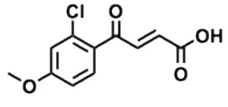

**Acid-059**

To a solution of 3-chloroanisole (1.5 g, 15.30 mmol) and maleic anhydride (2.0 g, 13.91 mmol) in DCM (50 mL) was added $AlCl_3$ (3.7 g, 27.81 mmol) portion wise. The mixture was stirred at room temperature for 24 hrs under $N_2$. The reaction mixture was quenched by ice-cooled 6N HCl (300 mL) and extracted with DCM (100 mL). The organic layer was washed with saturated $NaHCO_3$ aq. (300 mL). The water layer was adjusted to pH=3 with 1N HCl and extracted with DCM (100 mL x3). The combined organic layers were washed with brine (200 mL), dried over $Na_2SO_4$, filtered and concentrated to give crude product. The crude was purified by silical gel chromatography (EtOAc:Petroleum ether = 0 – 100%) to give Acid- 059 (1.4 g, 5.82 mmol, 41.8%) as a yellow solid.

$^1$H NMR (500 MHz, DMSO) δ 7.72 (d, *J* = 8.7 Hz, 1H), 7.38 (d, *J* = 15.7 Hz, 1H), 7.23 (d, *J* = 2.5 Hz, 1H), 7.11 (dd, *J* = 8.7, 2.5 Hz, 1H), 6.54 (d, *J* = 15.7 Hz, 1H), 3.92 (s, 3H).

$^{13}$C NMR (126 MHz, DMSO) δ 197.16, 191.43, 173.33, 167.05, 163.70, 162.72, 162.55, 138.03, 133.48, 133.06, 132.83, 132.66, 129.70, 128.33, 116.56, 116.33, 113.84, 113.40, 56.50, 56.39, 41.04, 40.57, 40.48, 40.40, 40.31, 40.23, 40.14, 40.07, 39.98, 39.90, 39.81, 39.64, 39.48.

HRMS (ESI): m/z calculated for $[C_{11}H_9ClO_4 + Na]^+$ = 263.0082, found 263.0088

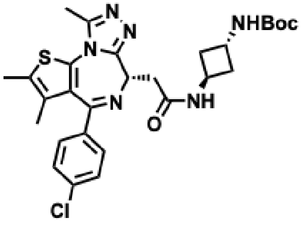

JQ1_PASW

Synthesized using General Procedure A, with (+)-JQ1-acid (100 mg, 0.25 mmol), tert-Butyl (trans-3-aminocyclobutyl)carbamate (022 mmol, 41.82 mg), HATU (142.3 mg, 0.374 mmol) and DIPEA (0.133 mL, 0.748 mmol). The crude residue was purified by silica gel chromatography (0 – 10% MeOH in DCM ) to afford 74 mg of the title compound (52.1%) as a brown solid.

$^1$H NMR (500 MHz, $CDCl_3$) δ 7.36 – 7.30 (m, 2H), 7.28 – 7.24 (m, 2H), 4.78 – 4.68 (m, 1H), 4.54 (dd, *J* = 7.6, 6.4 Hz, 1H), 4.37 (p, *J* = 7.2 Hz, 1H), 4.14 (s, 1H), 3.47 (dd, *J* = 14.3, 7.6 Hz, 1H), 3.32 – 3.21 (m, 1H), 2.60 (s, 3H), 2.36 – 2.29 (m, 4H), 2.28 – 2.23 (m, 2H), 2.19 (ddd, *J* = 12.7, 6.5, 3.6 Hz, 1H), 1.60 (s, 3H), 1.37 (s, 9H).

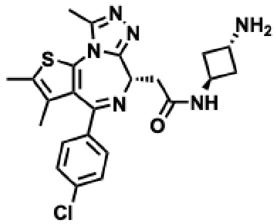

JQ1_PASW-$NH_2$

To a solution of **JQ1_PASW** (200 mg, 0.35 mmol, 1.0 eq.) in HCl/EtOAc (5 mL) and stirred at 25°C for 1 hr under $N_2$. The reaction mixture was concentrated under reduced pressure to give the title compound (200 mg, crude) as a yellow oil.

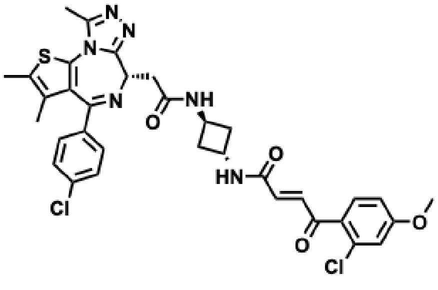

NVP-IKJ594

To a solution of **Acid-059** (74 mg, 0.30 mmol, 1.2 eq.), compound **JQ1_PASW-$NH_2$** (120 mg, 0.25 mmol, 1.0 eq.) and BOP (124 mg, 0.28, 1.1 eq.) in DMF (2 mL) was added DIPEA (0.221 mL, 1.28 mmol, 5.0 eq.). The mixture was stirred at room temperature overnight under $N_2$. The reaction mixture was quenched by 1N HCl (10 mL) and extracted with EtOAc (10 mL x 3). The combined organic layers were washed by brine (50 mL), dried over $Na_2SO_4$, filtered and concentrated under reduced pressure to give the residue. The crude product was purified by column (EtOAc:Petroleum ether = 0 – 100%) to afford the title compound as an off-white solid.

$^1$H NMR (500 MHz, $CDCl_3$) δ 7.36 – 7.30 (m, 2H), 7.28 – 7.24 (m, 2H), 4.78 – 4.68 (m, 1H), 4.54 (dd, *J* = 7.6, 6.4 Hz, 1H), 4.37 (p, *J* = 7.2 Hz, 1H), 4.14 (s, 1H), 3.47 (dd, *J* = 14.3, 7.6 Hz, 1H), 3.32 – 3.21 (m, 1H), 2.60 (s, 3H), 2.36 – 2.29 (m, 4H), 2.28 – 2.23 (m, 2H), 2.19 (ddd, *J* = 12.7, 6.5, 3.6 Hz, 1H), 1.60 (s, 3H), 1.37 (s, 9H).

$^{13}$C NMR (126 MHz, $CDCl_3$) δ 172.38, 166.09, 157.74, 157.29, 152.04, 138.96, 138.69, 134.26, 133.04, 132.98, 132.58, 131.93, 130.88, 79.42, 79.16, 78.91, 56.58, 55.96, 44.98, 44.20, 43.98, 41.36, 39.97, 39.63,

30.52, 20.76, 19.50, 16.51, 15.22, 14.17, 13.96.HRMS (ESI): m/z calculated for $[C_{34}H_{32}Cl_2N_6O_4S + H]^+$= 691.1656, found 691.1651.

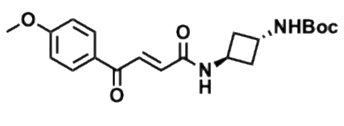

**EST1127**

Synthesized using General Procedure A, with (*E*)-4-(4-methoxyphenyl)-4-oxobut-2-enoic acid (0.890 g, 4.32 mmol), tert-Butyl (trans-3-aminocyclobutyl)carbamate (803.9 mg, 4.32 mmol), HATU (2.75 g, 8.63 mmol) and DIPEA (2.31 mL, 12.95 mmol). The crude residue was purified by silica gel chromatography (0 – 15% MeOH in DCM) to afford 312 mg of the title compound (19.3%) as a brown solid.

$^1$H NMR (500 MHz, $CDCl_3$) δ 8.08 – 8.01 (m, 2H), 7.94 (d, *J* = 15.1 Hz, 1H), 7.02 – 6.94 (m, 3H), 4.50 – 4.41 (m, 1H), 4.24 – 4.18 (m, 1H), 3.90 (s, 3H), 2.59 (s, 2H), 2.35 (h, *J* = 5.9 Hz, 4H), 1.44 (s, 9H).

$^{13}$C NMR (126 MHz, $CDCl_3$) δ 188.59, 164.51, 164.35, 155.74, 134.88, 134.83, 132.93, 131.47, 129.95, 114.20, 79.81, 77.40, 77.15, 76.90, 55.63, 49.98, 49.81, 49.63, 49.46, 49.29, 49.12, 48.95, 42.85, 41.85, 37.48, 37.46, 28.39.

HRMS (ESI): m/z calculated for $[C_{20}H_{26}N_2O_5 + Na]^+$ = 397.1734, found 397.1725

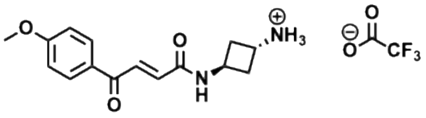

**AM-2-011**

Synthesized using General Procedure C.

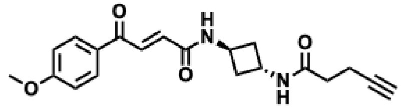

**AM-2-012**

Synthesized using General Procedure A, with **AM-2-011** (90 mg, 0.328 mmol), pent-4-ynoic acid (38.62 mg, 0.394 mmol), HATU (249.5 mg, 0.656 mmol), DIPEA (0.234 mL, 1.31 mmol). The crude residue was purified by silica gel chromatography (0-10% MeOH in DCM) to afford 42 mg of the title compound (36.12%) as a beige solid.

$^1$H NMR (500 MHz, DMSO) δ 8.93 (d, *J* = 7.2 Hz, 1H), 8.30 (d, *J* = 7.1 Hz, 1H), 8.04 – 7.96 (m, 2H), 7.75 (d, *J* = 15.3 Hz, 1H), 7.11 – 7.05 (m, 2H), 6.92 (d, *J* = 15.3 Hz, 1H), 4.35 (qd, *J* = 6.9, 1.1 Hz, 1H), 4.30 – 4.22 (m, 1H), 3.85 (s, 3H), 2.75 (t, *J* = 2.6 Hz, 1H), 2.37 – 2.30 (m, 2H), 2.27 – 2.18 (m, 6H).

$^{13}$C NMR (126 MHz, DMSO) δ 188.21, 170.25, 164.18, 163.47, 136.08, 132.28, 131.62, 129.99, 114.77, 84.21, 71.79, 56.14, 55.38, 41.82, 41.42, 40.57, 40.48, 40.41, 40.32, 40.24, 40.15, 40.07, 39.98, 39.81, 39.65, 39.48, 37.17, 34.59, 14.65.

HRMS (ESI): m/z calculated for $[C_{20}H_{22}N_2O_4 + H]^+$ = 355.1652, found 355.1644

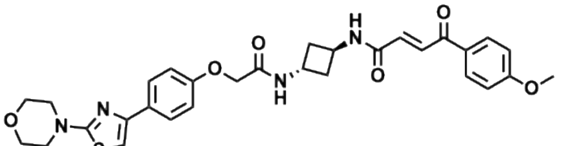

EST1140

Synthesized using General Procedure A, with **AM-2-011** (162.68 mg, 0.418 mmol), tert-butyl 2-(4-(2-morpholinothiazol-4-yl)phenoxy)acetate (122 mg, 0.381 mmol), HATU (289.6 mg, 0.761 mmol), DIPEA (0.339 mL, 1.90 mmol). The crude residue was purified by silica gel chromatography (0-10% MeOH in DCM) to afford 77.8 mg of the title compound (35.43%) as an off white solid.

$^1$H NMR (500 MHz, DMSO) δ 8.51 (d, *J* = 7.5 Hz, 1H), 8.35 (d, *J* = 7.1 Hz, 1H), 8.03 – 7.98 (m, 2H), 7.88 – 7.82 (m, 2H), 7.21 (s, 1H), 7.12 – 7.07 (m, 2H), 7.06 – 7.00 (m, 2H), 4.54 (s, 2H), 4.49 – 4.40 (m, 1H), 4.28 (tq, *J* = 7.6, 3.9 Hz, 1H), 3.90 (s, 3H), 3.82 – 3.76 (m, 4H), 3.49 (dd, *J* = 5.9, 3.8 Hz, 4H), 3.23 (t, *J* = 6.8 Hz, 2H), 2.49 (t, *J* = 6.7 Hz, 2H), 2.33 (ddd, *J* = 11.0, 8.2, 5.7 Hz, 2H), 2.23 (ddd, *J* = 12.5, 8.0, 5.0 Hz, 2H).

$^{13}$C NMR (101 MHz, DMSO-$D_6$) δ 188.29, 171.22, 167.65, 164.26, 163.59, 157.89, 150.84, 136.19, 132.34, 131.69, 130.08, 128.66, 127.53, 115.27, 114.85, 101.50, 67.55, 65.97, 56.20, 48.73, 41.77, 41.45, 36.99.

HRMS (ESI): calculated for $[C_{30}H_{32}N_4O_6S + H]^+$ = 577.2115, found 577.2109

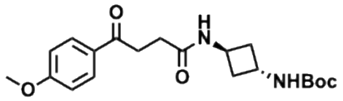

EL-1-013

Synthesized using General Procedure A, with 4-(4-methoxyphenyl)-4-oxobutanoic acid (100.0 mg, 0.480 mmol), tert-Butyl (trans-3-aminocyclobutyl)carbamate (98.40 mg, 0.528 mmol), HATU (365.24 mg, 0.961 mmol) and DIPEA (0.256 mL, 1.44 mmol). The crude residue was purified by silica gel chromatography (5 – 80% EtOAC in hexanes) to afford 33.20 mg of the title compound (18.36%) as a white solid.

$^1$H NMR (400 MHz, DMSO-D6) δ 8.16 (d, J = 7.0 Hz, 1H), 7.94 – 7.86 (m, 2H), 7.17 (d, J = 7.4 Hz, 1H), 7.03 – 6.96 (m, 2H), 4.08 (h, J = 6.6 Hz, 1H), 4.03 – 3.92 (m, 1H), 3.80 (s, 3H), 3.12 (t, J = 6.7 Hz, 2H), 2.38 (t, J = 6.7 Hz, 2H), 2.13 – 2.01 (m, 4H), 1.33 (s, 9H).

$^{13}$C NMR (101 MHz, DMSO-D6) δ 197.79, 171.27, 163.56, 155.34, 130.67, 114.37, 78.10, 56.04, 42.68, 41.10, 40.75, 40.70, 40.54, 40.49, 40.33, 40.28, 40.07, 39.86, 39.65, 39.45, 37.32, 33.47, 29.82, 28.78.

HRMS: calculated for $[C_{20}H_{28}N_2O_5 + H]^+$ = 376.1998, found 376.1987

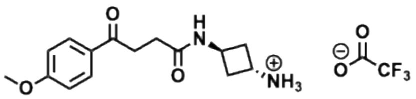

EL-1-014

Synthesized using General Procedure C.

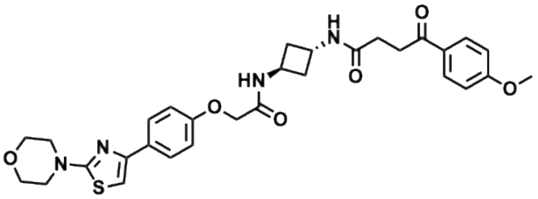

AM-2-040

Synthesized using General Procedure A, with **EL-1-014** (22.63 mg, 0.045 mmol), 4-(4-methoxyphenyl)-4-oxobutanoic acid (10.31 mg, 0.050 mmol), HATU (51.37 mg, 0.135 mmol), DIPEA (0.032 mL, 0.181 mmol). The crude residue was purified by silica gel chromatography (0-10% MeOH in DCM) to afford 14.2 mg of the title compound (54.5%) as an orange solid.

$^{1}$H NMR (500 MHz, DMSO) δ 8.51 (d, *J* = 7.5 Hz, 1H), 8.35 (d, *J* = 7.1 Hz, 1H), 8.03 – 7.98 (m, 2H), 7.88 – 7.82 (m, 2H), 7.21 (s, 1H), 7.12 – 7.07 (m, 2H), 7.06 – 7.00 (m, 2H), 4.54 (s, 2H), 4.49 – 4.40 (m, 1H), 4.28 (tq, *J* = 7.6, 3.9 Hz, 1H), 3.90 (s, 3H), 3.82 – 3.76 (m, 4H), 3.49 (dd, *J* = 5.9, 3.8 Hz, 4H), 3.23 (t, *J* = 6.8 Hz, 2H), 2.49 (t, *J* = 6.7 Hz, 2H), 2.33 (ddd, *J* = 11.0, 8.2, 5.7 Hz, 2H), 2.23 (ddd, *J* = 12.5, 8.0, 5.0 Hz, 2H).

$^{13}$C NMR (126 MHz, DMSO) δ 197.71, 171.28, 171.15, 167.51, 163.49, 157.81, 150.76, 130.61, 130.02, 128.56, 127.44, 115.17, 115.02, 114.30, 101.51, 101.42, 67.44, 65.90, 55.98, 48.65, 41.35, 41.30, 41.26, 41.21, 40.57, 40.48, 40.41, 40.32, 40.24, 40.15, 40.07, 39.98, 39.90, 39.82, 39.65, 39.48, 37.09, 37.03, 33.41, 29.76.

HRMS (ESI): calculated for $[C_{30}H_{34}N_4O_6S + H]^+$ = 579.2272, found 579.2264

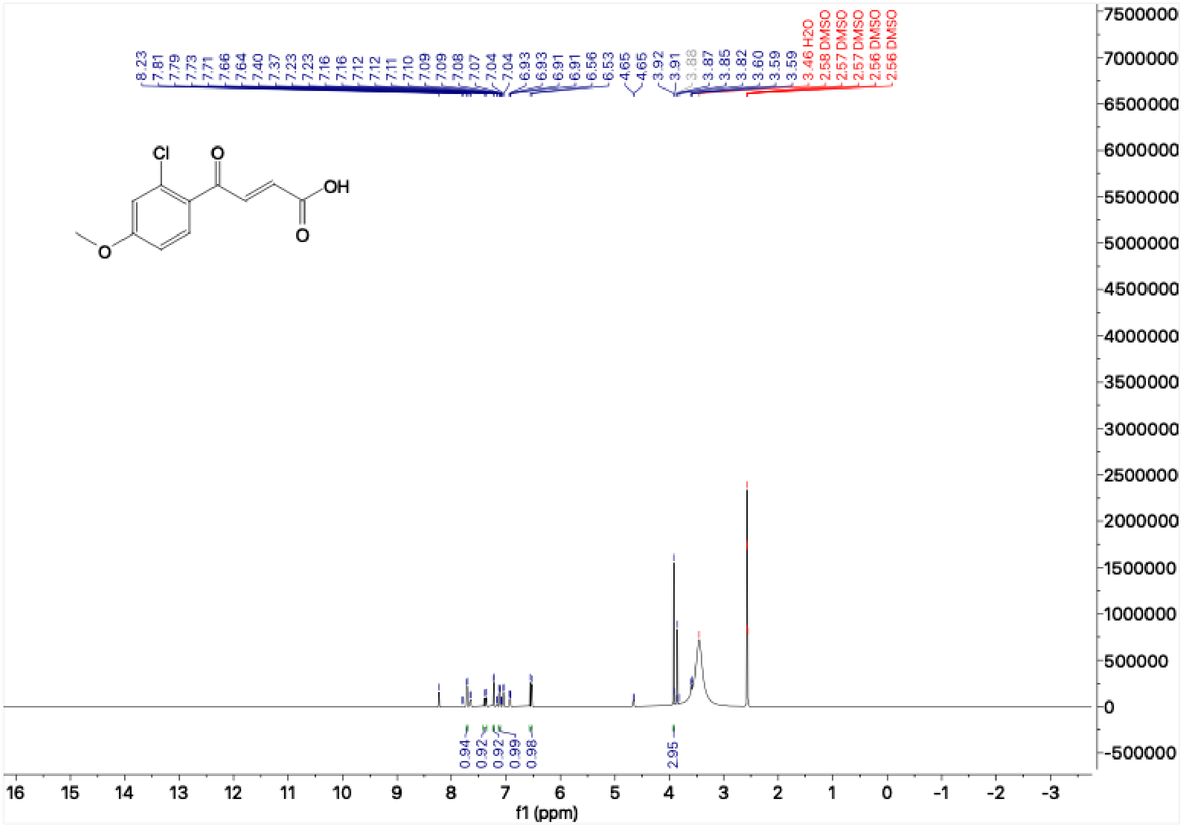

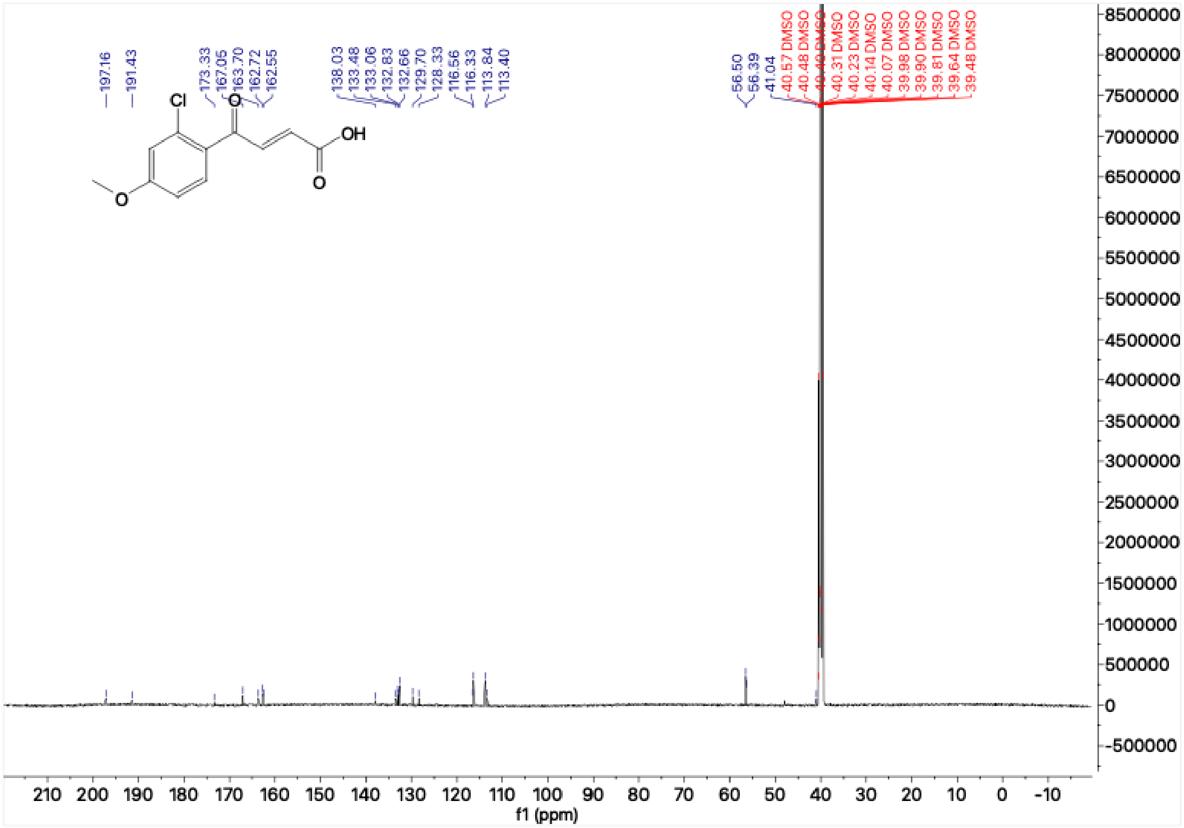

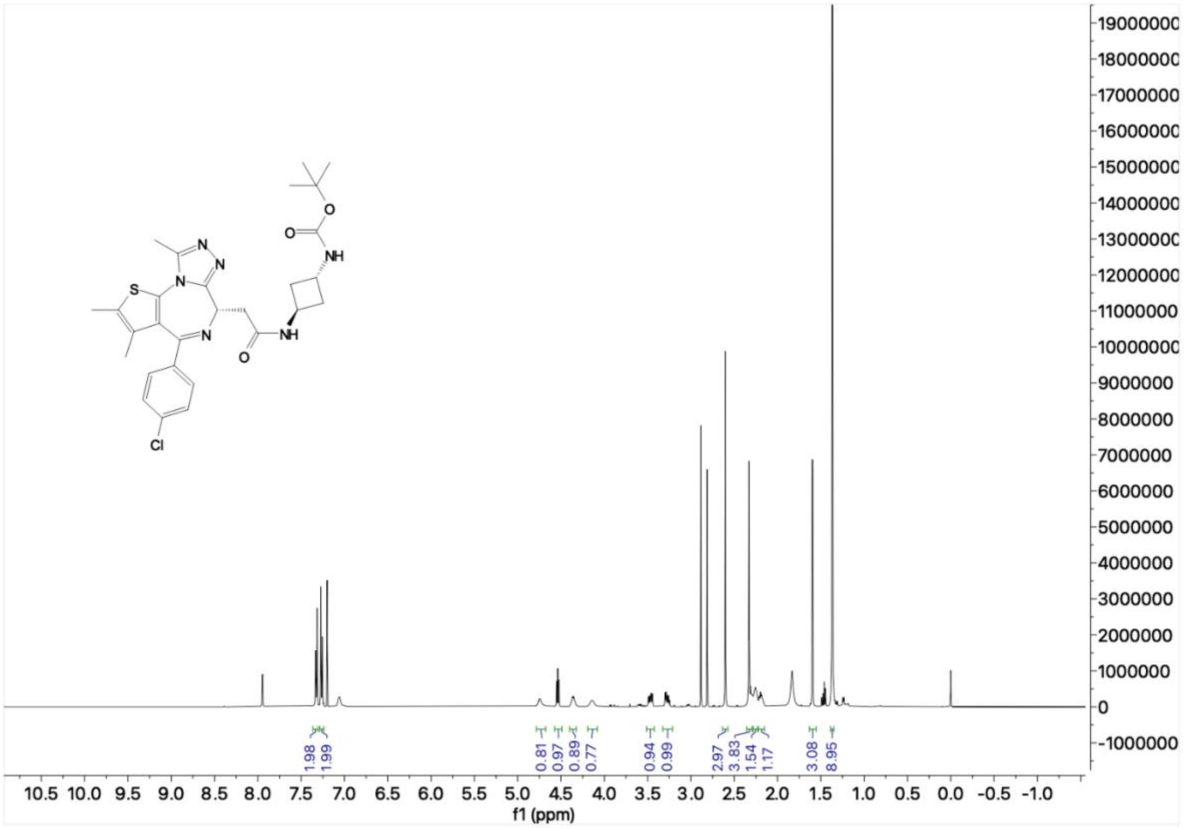

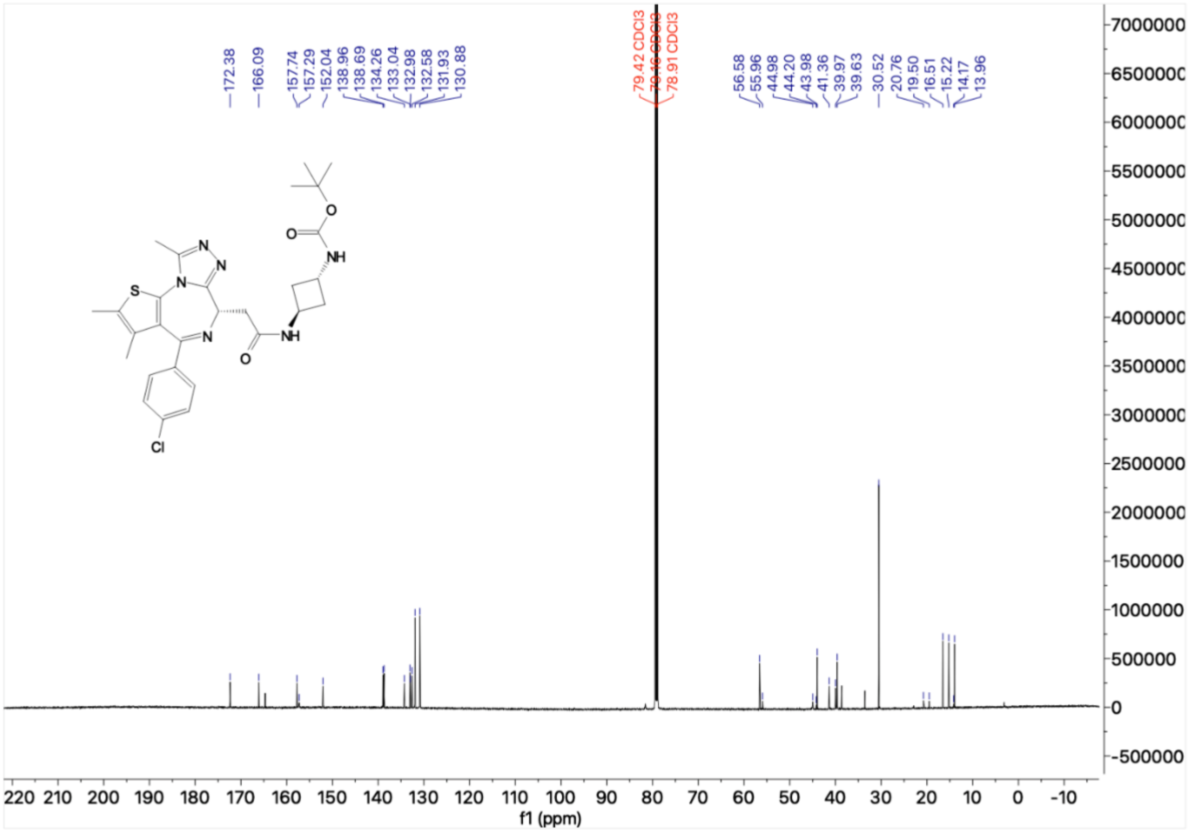

172.38
166.09
157.74
157.29
152.04
138.96
138.69
134.26
133.04
132.98
132.58
131.93
130.88
79.42 CDCl3
78.91 CDCl3
56.58
55.96
44.98
44.20
43.98
41.36
39.97
39.63
30.52
20.76
19.50
16.51
15.22
14.17
13.96
S
N
NH
O
Cl
f1 (ppm)


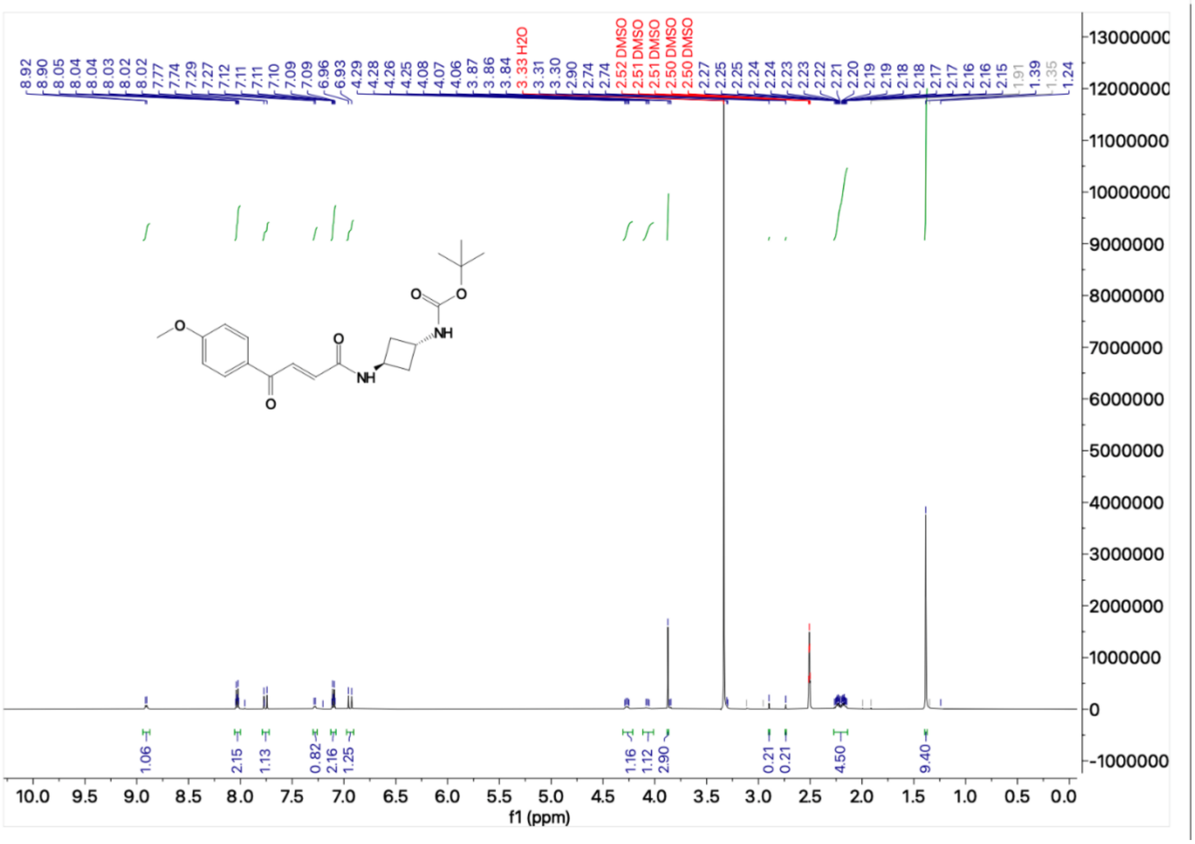

8.92
8.90
8.05
8.04
8.03
8.02
8.02
7.77
7.74
7.29
7.27
7.12
7.11
7.11
7.10
7.09
7.09
6.96
6.93
4.29
4.28
4.26
4.25
4.08
4.07
4.06
3.87
3.86
3.84
3.33 H2O
3.31
3.30
2.90
2.74
2.74
2.52 DMSO
2.51 DMSO
2.51 DMSO
2.50 DMSO
2.50 DMSO
2.27
2.25
2.25
2.24
2.24
2.23
2.23
2.22
2.21
2.20
2.19
2.19
2.18
2.18
2.17
2.17
2.16
2.16
2.15
1.91
1.39
1.35
1.24
O
NH
1.06
2.15
1.13
0.82
2.16
1.25
1.16
1.12
2.90
0.21
0.21
4.50
9.40
f1 (ppm)

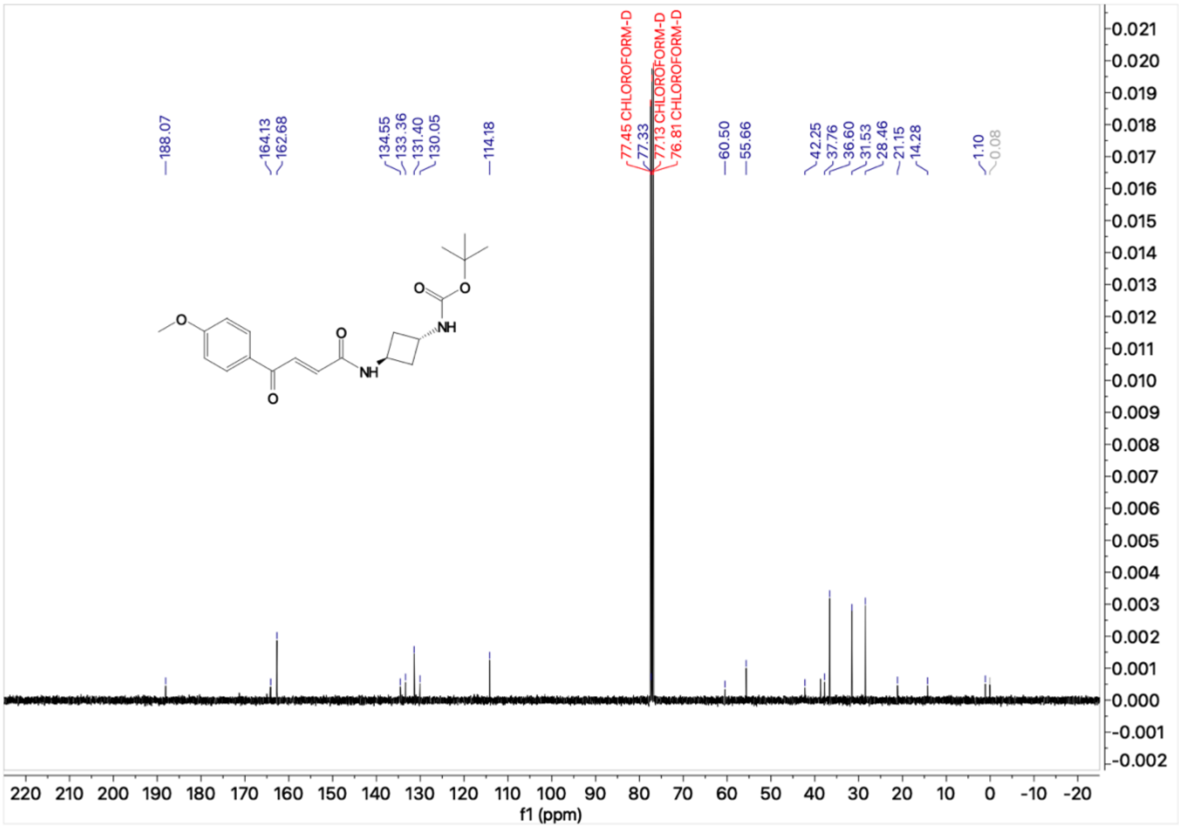
188.07
164.13
162.68
134.55
133.36
131.40
130.05
114.18
77.45 CHLOROFORM-D
77.33
77.13 CHLOROFORM-D
76.81 CHLOROFORM-D
60.50
55.66
42.25
37.76
36.60
31.53
28.46
21.15
14.28
1.10
-0.08
O
O
O
O
NH
NH
O
O
220 210 200 190 180 170 160 150 140 130 120 110 100 90 80 70 60 50 40 30 20 10 0 -10 -20
f1 (ppm)
0.021
0.020
0.019
0.018
0.017
0.016
0.015
0.014
0.013
0.012
0.011
0.010
0.009
0.008
0.007
0.006
0.005
0.004
0.003
0.002
0.001
0.000
-0.001
-0.002

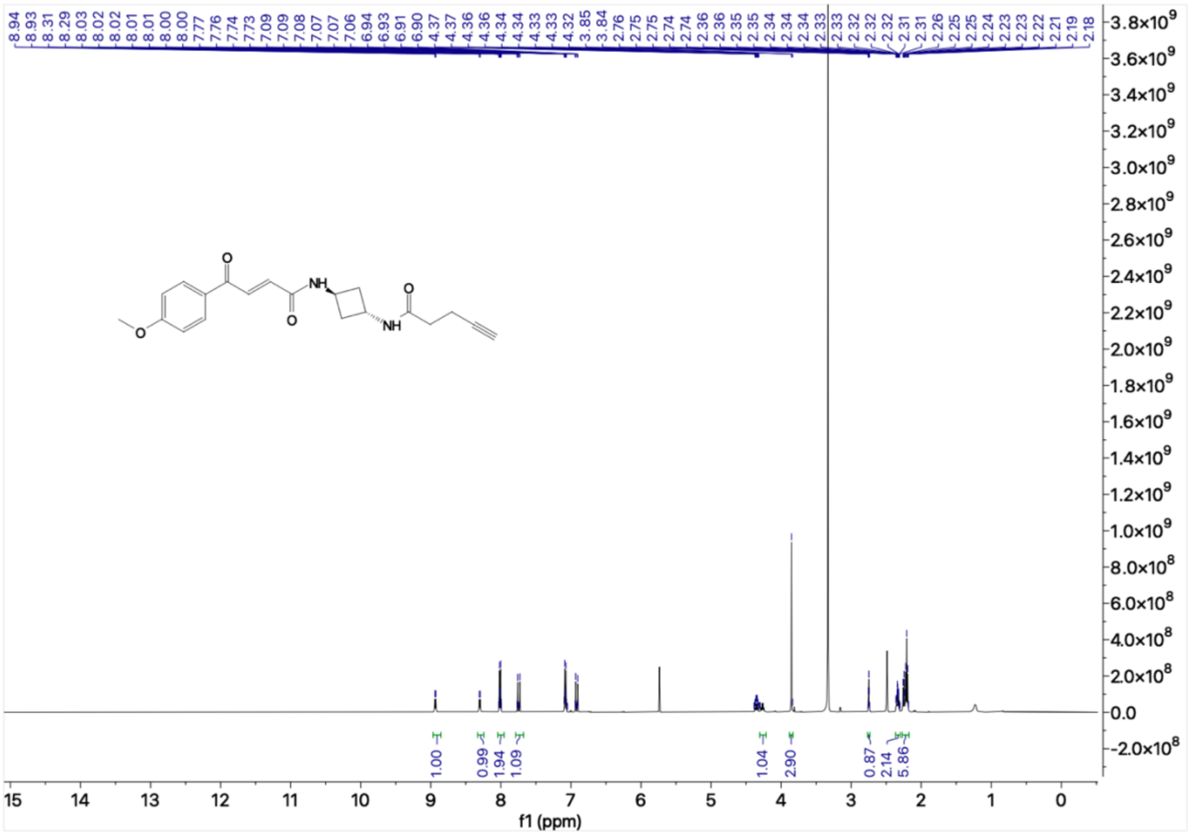
8.94 8.93 8.31 8.29 8.03 8.02 8.02 8.01 8.00 7.77 7.76 7.74 7.73 7.09 7.09 7.08 7.07 7.07 7.06 6.94 6.93 6.91 6.90 4.37 4.37 4.36 4.36 4.34 4.34 4.33 4.33 4.32 3.85 3.84 2.76 2.75 2.75 2.74 2.74 2.36 2.36 2.35 2.35 2.34 2.34 2.34 2.33 2.33 2.32 2.32 2.32 2.32 2.31 2.26 2.25 2.24 2.23 2.23 2.22 2.21 2.19 2.18
O
NH
O
O
NH
O
1.00
0.99
1.94
1.09
1.04
2.90
0.87
2.14
5.86
f1 (ppm)

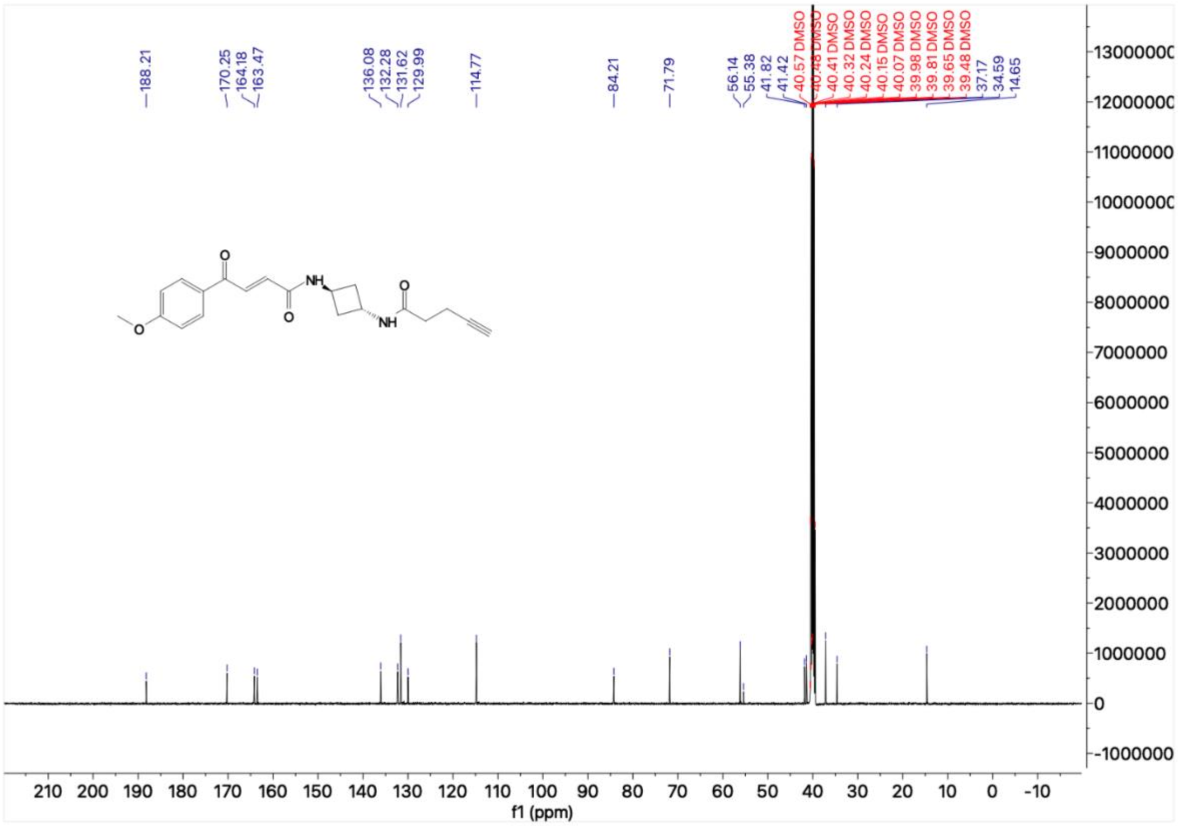
188.21
170.25
164.18
163.47
136.08
132.28
131.62
129.99
114.77
84.21
71.79
56.14
55.38
41.82
41.42
40.57 DMSO
40.48 DMSO
40.41 DMSO
40.32 DMSO
40.24 DMSO
40.15 DMSO
40.07 DMSO
39.98 DMSO
39.81 DMSO
39.65 DMSO
39.48 DMSO
37.17
34.59
14.65
O
NH
O
O
NH
O
f1 (ppm)

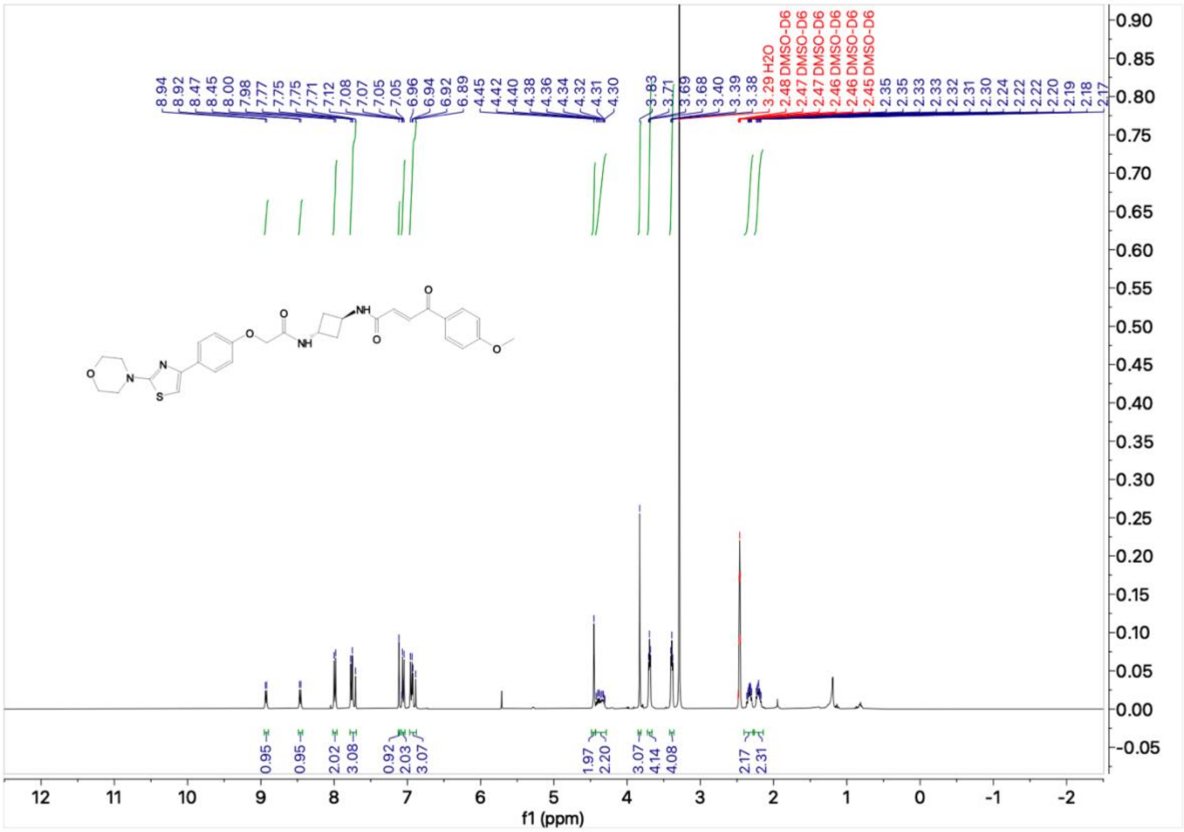
8.94 8.92 8.47 8.45 8.00 7.98 7.77 7.75 7.75 7.71 7.12 7.08 7.07 7.05 7.05 6.96 6.94 6.92 6.89 4.45 4.42 4.40 4.38 4.36 4.34 4.32 4.31 4.30 3.83 3.71 3.69 3.68 3.40 3.39 3.38
3.29 H2O
2.48 DMSO-D6
2.47 DMSO-D6
2.47 DMSO-D6
2.46 DMSO-D6
2.46 DMSO-D6
2.45 DMSO-D6
2.35 2.35 2.33 2.33 2.32 2.31 2.30 2.24 2.22 2.22 2.20 2.19 2.18 2.17
0.95 0.95 2.02 3.08 0.92 2.03 3.07 1.97 2.20 3.07 4.14 4.08 2.17 2.31
f1 (ppm)

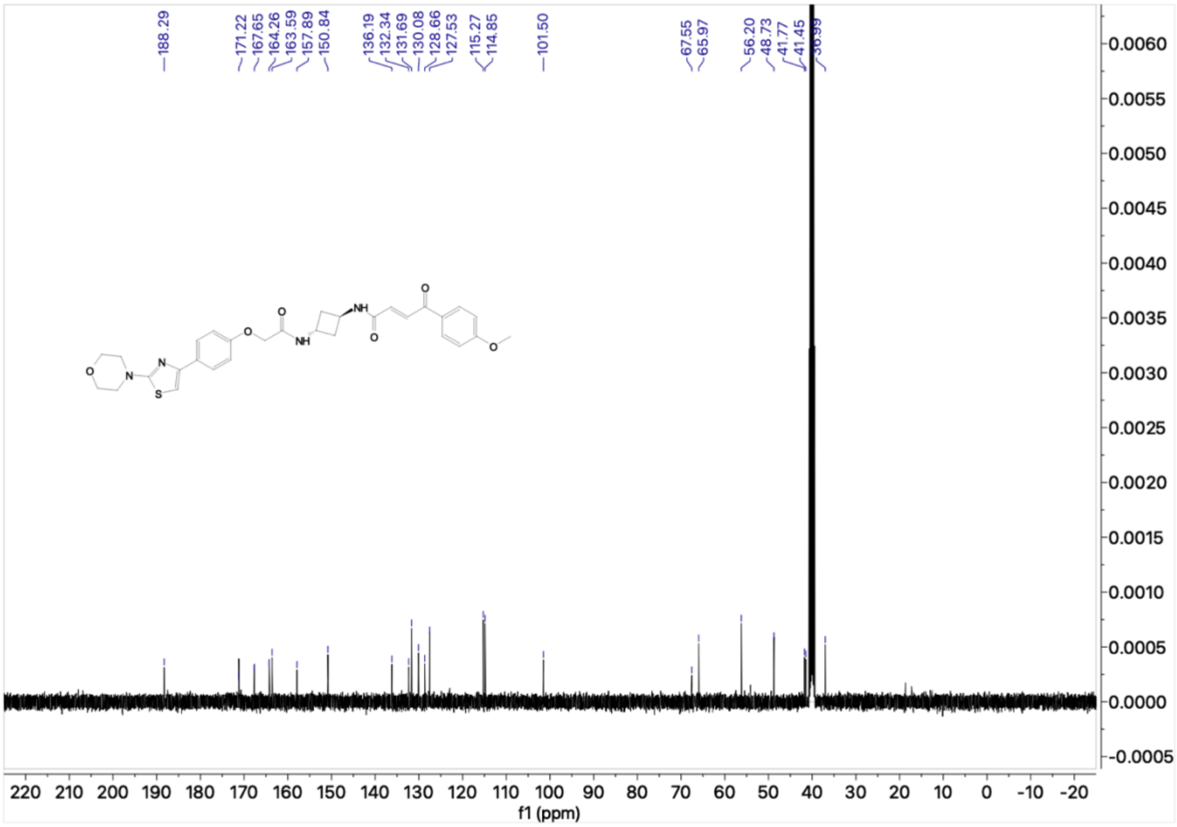
188.29
171.22 167.65 164.26 163.59 157.89 150.84
136.19 132.34 131.69 130.08 128.66 127.53 115.27 114.85
101.50
67.55 65.97
56.20 48.73 41.77 41.45 36.99
f1 (ppm)

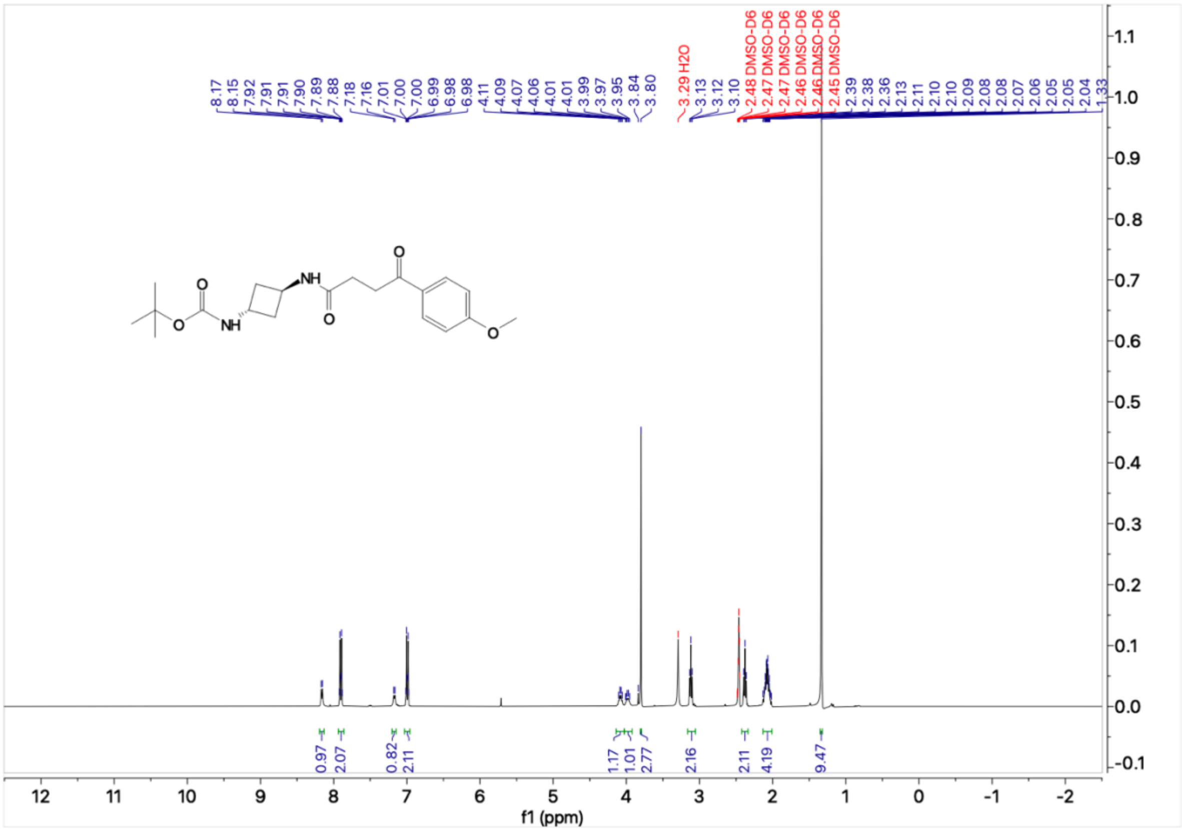
8.17 8.15 7.92 7.91 7.91 7.90 7.89 7.88 7.18 7.16 7.01 7.00 7.00 6.99 6.98 6.98 4.11 4.09 4.07 4.06 4.01 4.01 3.99 3.97 3.95 3.84 3.80
3.29 H2O
3.13 3.12 3.10
2.48 DMSO-D6 2.47 DMSO-D6 2.47 DMSO-D6 2.46 DMSO-D6 2.46 DMSO-D6 2.45 DMSO-D6
2.39 2.38 2.36 2.13 2.11 2.10 2.10 2.09 2.08 2.08 2.07 2.06 2.05 2.05 2.04 1.33
0.97 2.07 0.82 2.11 1.17 1.01 2.77 2.16 2.11 4.19 9.47
f1 (ppm)

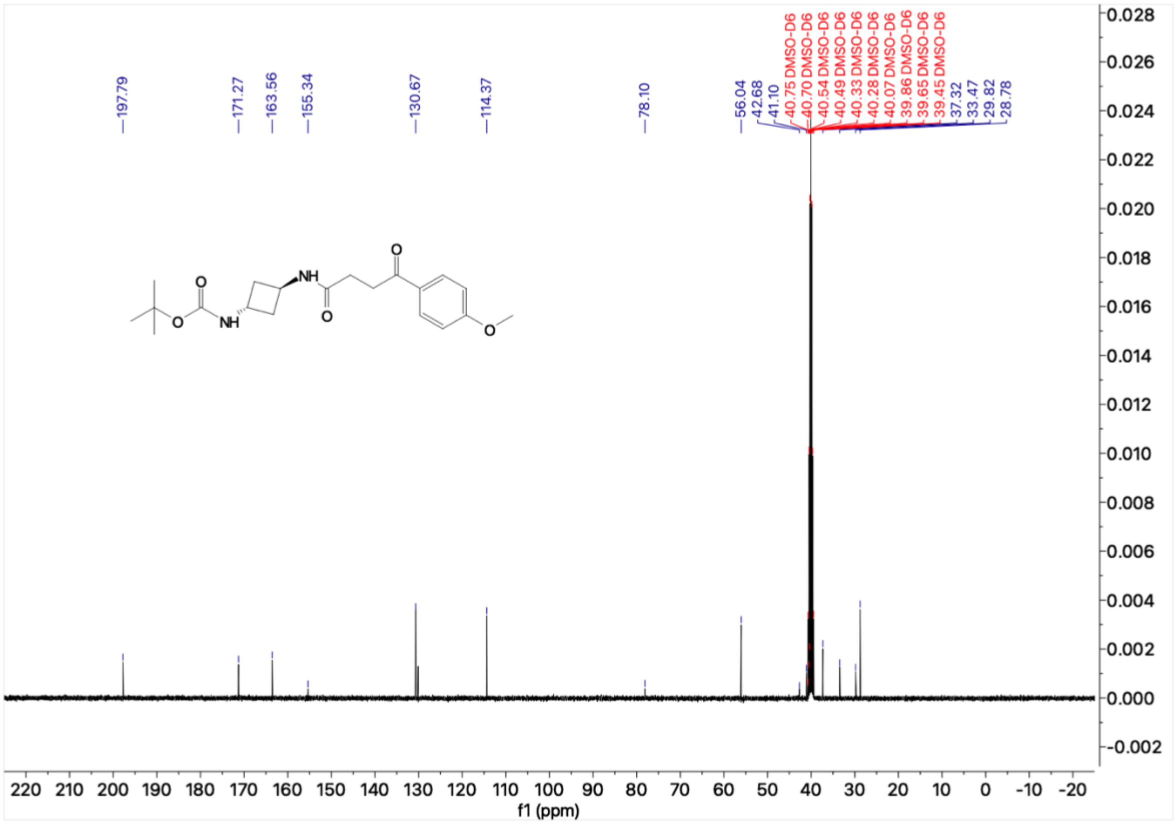
197.79 171.27 163.56 155.34 130.67 114.37 78.10 56.04 42.68 41.10
40.75 DMSO-D6 40.70 DMSO-D6 40.54 DMSO-D6 40.49 DMSO-D6 40.33 DMSO-D6 40.28 DMSO-D6 40.07 DMSO-D6 39.86 DMSO-D6 39.65 DMSO-D6 39.45 DMSO-D6
37.32 33.47 29.82 28.78
f1 (ppm)

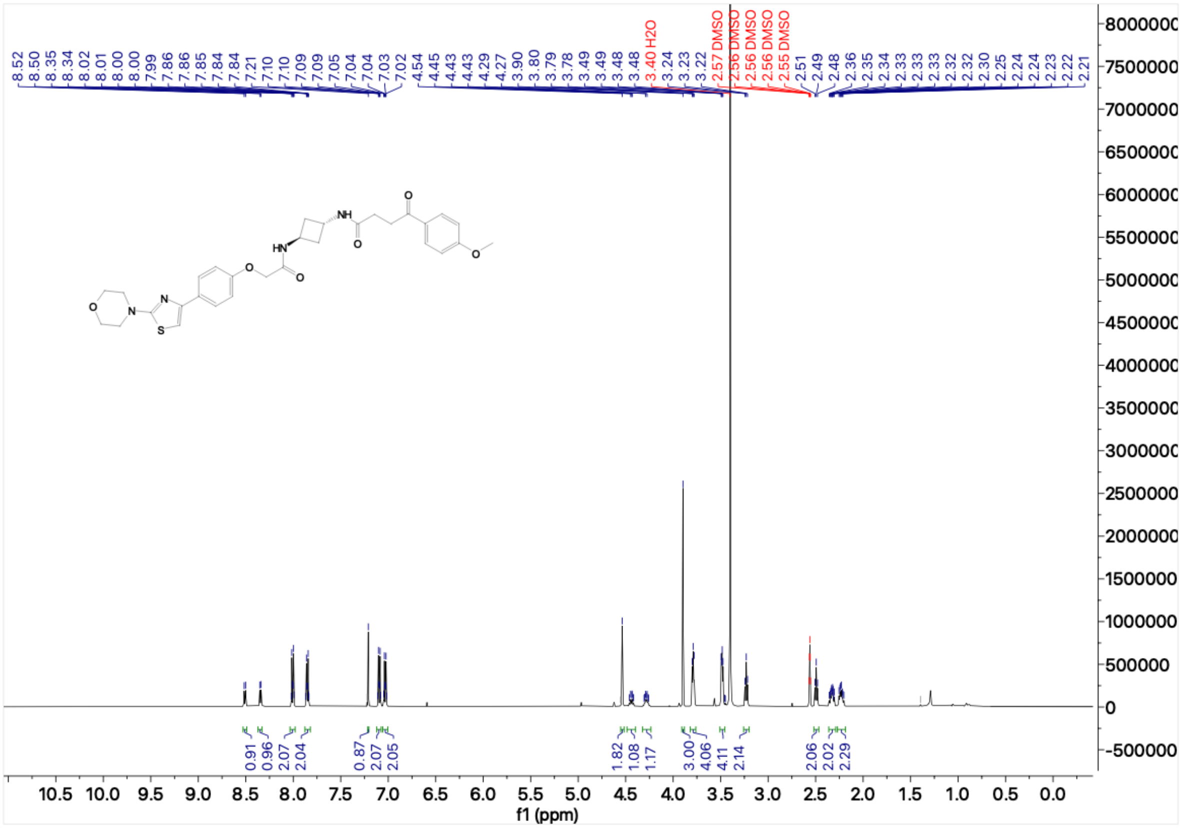

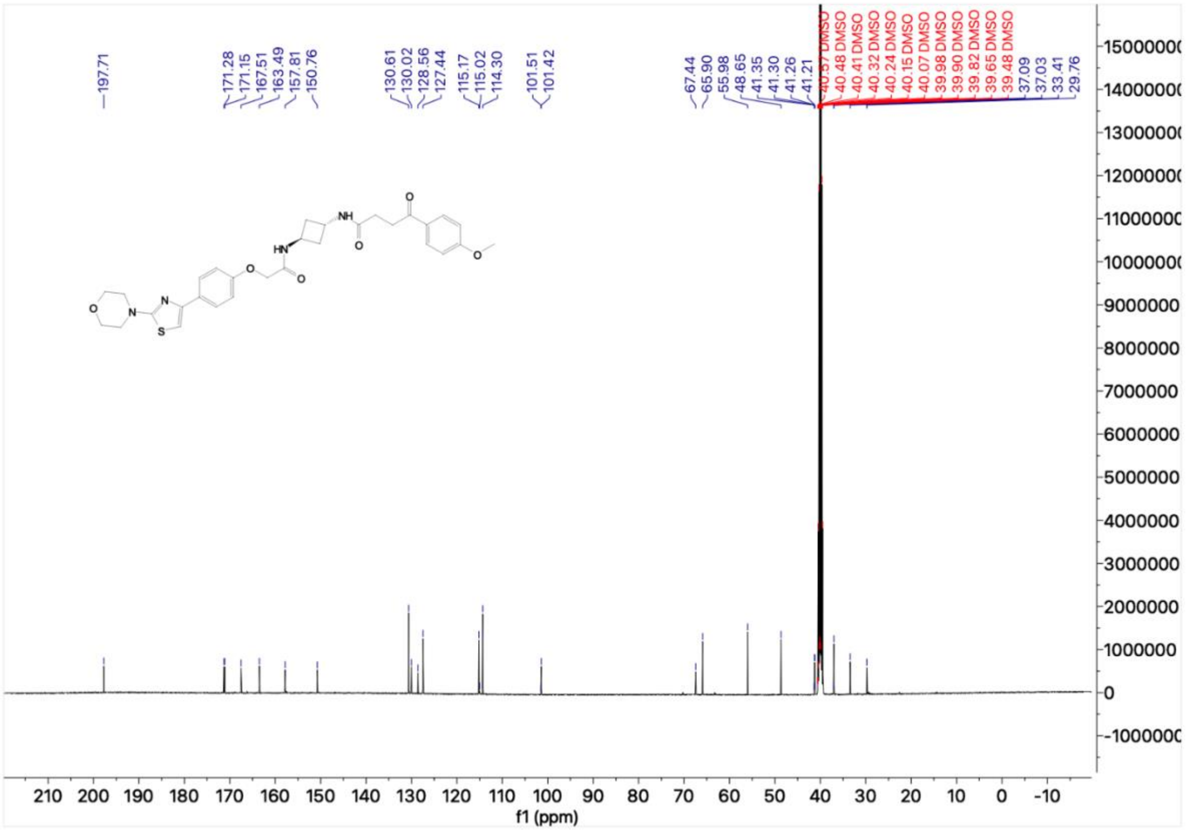

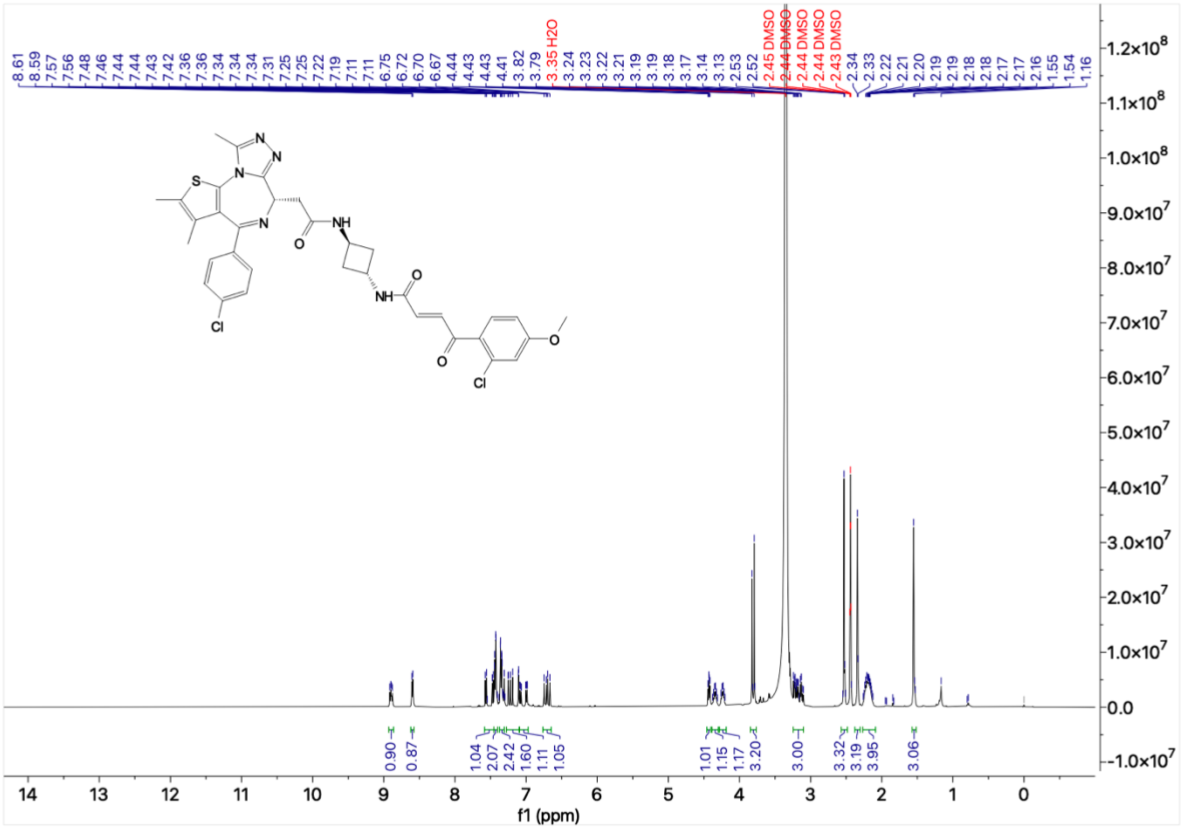
8.61 8.59 7.57 7.56 7.48 7.46 7.44 7.44 7.43 7.42 7.36 7.36 7.34 7.34 7.34 7.31 7.25 7.25 7.22 7.19 7.11 7.11 6.75 6.72 6.70 6.67 4.44 4.43 4.43 4.41 3.82 3.79 3.35 H2O 3.24 3.23 3.22 3.21 3.19 3.19 3.18 3.17 3.14 3.13 2.53 2.52 2.45 DMSO 2.44 DMSO 2.44 DMSO 2.44 DMSO 2.43 DMSO 2.34 2.33 2.22 2.21 2.20 2.19 2.19 2.18 2.18 2.17 2.17 2.16 1.55 1.54 1.16
0.90 0.87 1.04 2.07 2.42 1.60 1.11 1.05 1.01 1.15 1.17 3.20 3.00 3.32 3.19 3.95 3.06
f1 (ppm)

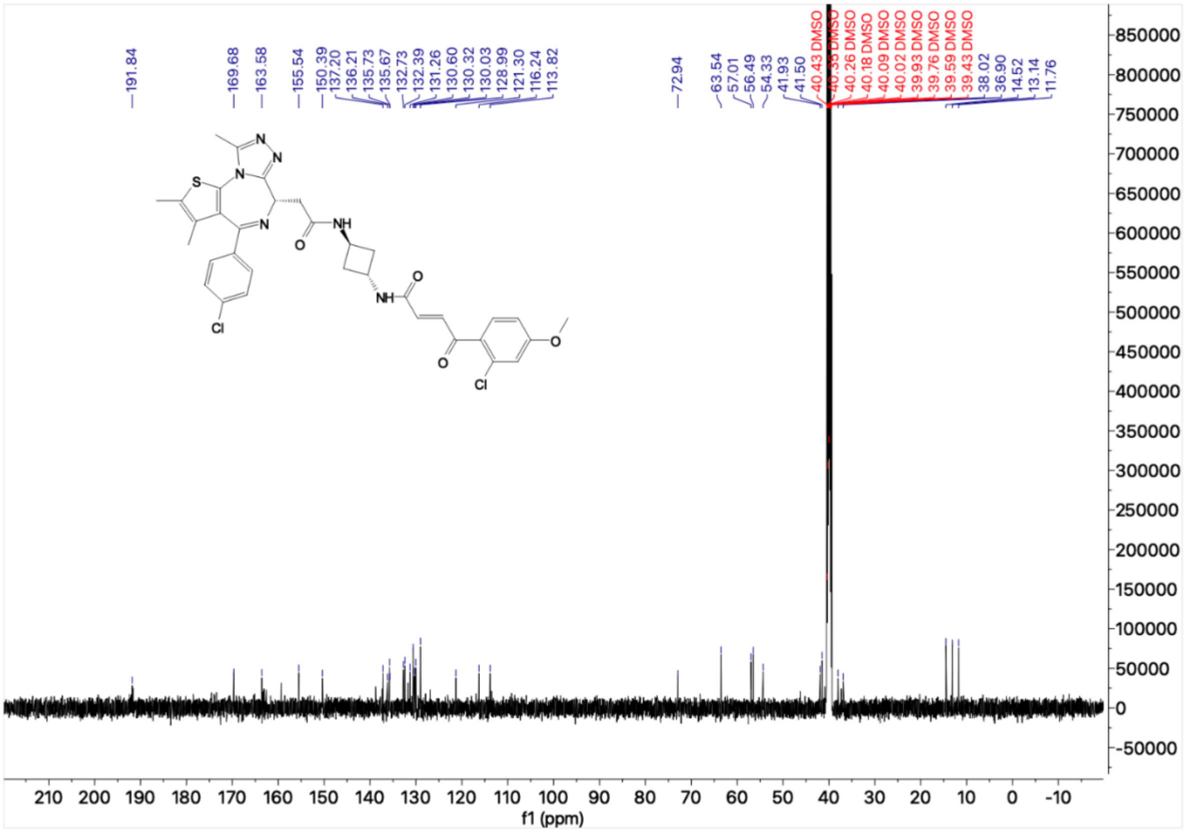
191.84 169.68 163.58 155.54 150.39 137.20 136.21 135.73 135.67 132.73 132.39 131.26 130.60 130.32 130.03 128.99 121.30 116.24 113.82 72.94 63.54 57.01 56.49 54.33 41.93 41.50 40.43 DMSO 40.35 DMSO 40.26 DMSO 40.18 DMSO 40.09 DMSO 40.02 DMSO 39.93 DMSO 39.76 DMSO 39.59 DMSO 39.43 DMSO 38.02 36.90 14.52 13.14 11.76
f1 (ppm)
